# Secondary motifs enable concentration-dependent regulation by Rbfox family proteins

**DOI:** 10.1101/840272

**Authors:** Bridget E. Begg, Marvin Jens, Peter Y. Wang, Christopher B. Burge

## Abstract

The Rbfox family of splicing factors regulate alternative splicing during animal development and in disease, impacting thousands of exons in the maturing brain, heart, and muscle. Rbfox proteins have long been known to bind to the RNA sequence GCAUG with high affinity, but just half of Rbfox CLIP peaks contain a GCAUG motif. We incubated recombinant RBFOX2 with over 60,000 transcriptomic sequences to reveal significant binding to several moderate-affinity, non-GCAYG sites at a physiologically relevant range of RBFOX concentrations. We find that many of these “secondary motifs” bind Rbfox robustly *in vivo* and that several together can exert regulation comparable to a GCAUG in a trichromatic splicing reporter assay. Furthermore, secondary motifs regulate RNA splicing in neuronal development and in neuronal subtypes where cellular Rbfox concentrations are highest, enabling a second wave of splicing changes as Rbfox levels increase.

## Introduction

The conserved Rbfox family of RNA-binding proteins (RBPs), which play critical roles in animal development, have been the subjects of numerous genetic, biochemical and structural studies since their discovery over twenty-five years ago. *Rbfox1* was identified in 1994 as the genomic locus *feminizing on X* (*fox-1*) in *C. elegans*; it was subsequently shown that the encoded protein peaks in expression in larval developmental stages and controls sex determination by repressing the dosage compensation factor *xol-1* post-transcriptionally^1, 2^. Mammalian *Rbfox1 (A2BP1)* and its paralogs *Rbfox2 (RBM9)* and *Rbfox3 (NeuN)* are highly expressed in heart, skeletal muscle, and in the adult brain, with a characteristic spike in expression in late neuronal development^3–6^. While Rbfox proteins are predominantly nuclear, where they regulate pre-mRNA splicing, some isoforms are also expressed in the cytoplasm, where they regulate RNA stability^7^.

The three mammalian paralogs have high sequence identity (94-99%) in their RNA-binding domains (RBDs), substantially overlapping gene targets, and similar activities, making single knockouts difficult to interpret and regulatory targets challenging to define^8, 9^. Nonetheless, mouse *Rbfox1* central nervous system (CNS) knockouts have seizures and defects in neuronal excitability, while *Rbfox2* CNS knockout mice exhibit dramatic defects in cerebellar development^9, 10^. Binding targets of RBFOX1, both mRNA splicing- and mRNA stability-related, were found to be enriched for genes related to autism, axon and dendrite formation, and electrophysiology in mice^11–14^, and mutations in *Rbfox1* and *Rbfox3* have been associated with autism and epilepsy in humans^15–18^. Recently, it was shown that Rbfox triple knockout mouse embryonic stem cells fail to develop a mature splicing profile when differentiated into ventral motor neurons, underscoring the roles of these proteins in proper neuronal maturation^8^.

In 2003, RBFOX1 was found to bind the pentanucleotide GCAUG with high affinity to regulate alterative splicing through binding to flanking intronic regions^19^. (U)GCAUG had been established as a dominant signal in computational analyses of neuronal alternative splicing, and Rbfox proteins have since been shown to regulate numerous alternative exons through this motif in a variety of neuronal subtypes^8, 14, 20, 21^. Binding to the GCAUG motif is mediated by canonical and non-canonical interactions between the protein’s single RNA recognition motif (RRM) and the RNA^22^. Generally, binding of Rbfox proteins to motifs in the proximal ∼200 nucleotides (nt) of the intron downstream of an alternative exon activates exon inclusion, while binding in the proximal upstream intron or alternative exon promotes exon exclusion^23^. However, more distal motifs are also able to regulate splicing through RNA bridge formation^24^. Proteins recruited by the Ala/Tyr/Gly-rich C-terminal domain of Rbfox mediate its effects on splicing, whereas its effects on stability may be related to competition with other RBPs and microRNAs^13, 25^.

Although Rbfox binding to canonical GCAUG motifs and rarer GCACG motifs is well characterized, studies of Rbfox binding sites *in vivo* using crosslinking-immunoprecipitation (CLIP) have observed that about half of CLIP peaks lack an associated GCAYG (Y = C or U) motif^23^, suggesting the existence of additional binding determinants^26^. Indeed, recent work has observed GUGUG motifs and motifs differing by one base from the canonical GCAUG in CLIP peaks^12, 27–29^, and RBFOX1 was shown to compete with MBNL1 at a GCCUG motif^30^. It has been proposed that binding to GUGUG is mediated by partner proteins such as SUP12 or members of the large assembly of splicing regulators (LASR) complex^31–33^. Direct binding of Rbfox proteins to these motifs has not been excluded, but if such binding occurs it is not known to have any biological consequence. Almost all studies of Rbfox binding to date have filtered their datasets for the presence of a GCAUG element, discarding many potential binding sites^8–10, 12–14, 34^.

Studies of Rbfox *in vivo* binding and regulation using CLIP have identified important regulatory targets and uncovered complex regulatory networks. However, CLIP is known to have substantial rates of false-positives and false-negatives^35–37^, making it challenging to confidently infer binding sites and motifs. Here we employ RNA Bind-n-Seq with natural sequences (nsRBNS) as a biochemical approach to refine our understanding of Rbfox binding across mammalian transcriptomes^28, 38^. Our method detects strong binding to the canonical GCAYG Rbfox motif, but also detects moderate binding to several additional motif variants. We show that Rbfox proteins bind to these secondary motifs *in vivo*, and exert regulatory activity in a manner dependent on Rbfox concentration. Furthermore, we find evidence of important roles for secondary motifs in splicing programs involved in neuronal differentiation and subtype specification, and we propose a model for Rbfox regulation that incorporates these motifs.

## Results

### Natural sequence RBNS recovers known features of RBFOX2 binding

To better understand the transcriptomic RNA binding preferences of RBFOX2, we designed nsRBNS library based on naturally-occurring mammalian 3’ untranslated region (UTR) sequences (Figure 1a). To construct the library, ∼2,200 well-annotated human and orthologous mouse 3’ UTRs from three groups of genes were selected and confirmed to match the base and *5*mer composition of the entire 3’ UTR transcriptome (Figure 1b). The resulting library of 64,319 oligonucleotides represents ∼10% of the 3’ UTR transcriptome of human HepG2 cells (data not shown). The 3’ UTRs of each of these transcripts were then divided into overlapping 110-base segments (Supplementary Table 1). These sequences occur naturally and are much longer than those used in random sequence RBNS (20-40 nt), enabling analysis of binding of RBFOX2 to motif clusters as well as individual motifs.

**Figure 1.**
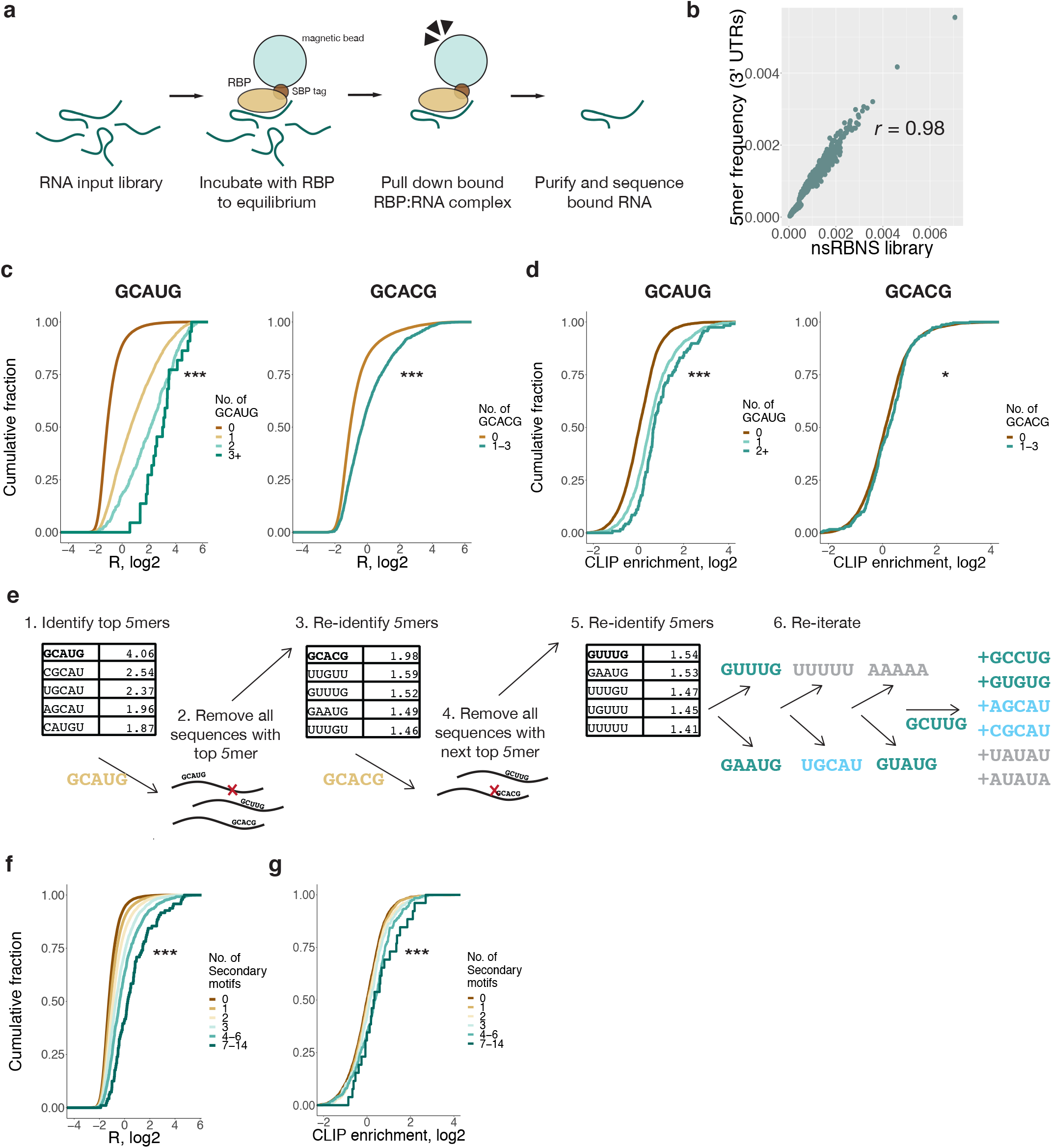
3’ UTR natural sequence nsRBNS (nsRBNS) with RBFOX2 captures variation in binding affinity arising from conserved elements in natural RNA sequences. a. Schematic of nsRBNS. Recombinant protein is incubated with a designed RNA library to equilibrium and bound RBP:RNA complexes are pulled down via a streptavidin-coated magnetic bead. Oligonucleotides are then sequenced and the enrichment (*R*) value of each is calculated as their frequency in the pulldown library divided by their frequency in the input library. b. nsRBNS 5mer frequencies are highly correlated with 5mer frequencies of the 3’ UTR transcriptome. Pearson correlation. c. nsRBNS sequences containing Rbfox primary motifs GCAUG and GCACG increase in *R* ((reads per million)input/(reads per million)pulldown) with increased motif count. Significance shown between lowest and highest counts was assessed by Wilcoxon Rank-Sum test (*** P < 0.001). d. Increased RBFOX2 eCLIP in HepG2 cells with increased motif count in transcriptomic regions corresponding to those in nsRBNS library normalized to IgG control. Significance shown between lowest and highest counts by Wilcoxon Rank-Sum test (*** P < 0.001, * P < 0.05). e. An iterative method discovered moderate binding by RBFOX2 to six additional motifs of the sequence format GNNUG in nsRBNS. After identifying the top enriched pentamer, all sequences containing that pentamer were removed for a subsequent round of enrichment analysis to reveal other enriched pentamers. After four rounds of enrichment analysis, remaining *5*mers of the format GNNUG were also included as secondary motifs f. nsRBNS sequences containing increasing numbers of secondary motifs have corresponding increases in *R* value. Significance shown between lowest and highest counts by Wilcoxon Rank-Sum test (*** P < 0.001). g. RBFOX2 eCLIP in HepG2 cells shows increased RBFOX2 peak enrichment with increased secondary motif count at library positions in the transcriptome. RBFOX2 peaks were compared to an IgG control to determine enrichments. Significance shown between lowest and highest counts by Wilcoxon Rank-Sum (*** P < 0.001).

Using a range of protein concentrations in an RBNS experiment allows detection of binding sites in different affinity ranges^28^. We therefore performed nsRBNS with recombinant, tagged RBFOX2 at 4 nM, 14 nM, 43 nM, 121 nM (in technical duplicate), 365 nM and 1.1µM, as well as a 0 nM (no protein) control, with a constant 250 nM total RNA concentration. This range, spanning 2.5 orders of magnitude, is comparable to the large range of natural total Rbfox family expression, which varies in RNA expression by at least three orders of magnitude across different cell types^4^, and by at least an order of magnitude during neuronal differentiation^39^. The range of RBFOX2 concentration tested here captures binding to sequences that are bound and active *in vivo*, and overlaps substantially with the physiological range of Rbfox expression (see below). Enrichment (*R*) values were calculated for each individual oligonucleotide as the ratio of the frequency in the bound pool of RNA over the frequency in control (no protein) conditions (Supplementary Table 2). *R* values of bound oligonucleotides at similar concentrations were strongly correlated (*r* ≥ 0.74) (Supplementary Figure 1a), indicating high reproducibility of oligonucleotide-specific binding. *R* values of GCAUG were highest at the 365 nM concentration, as observed in random sequence RBNS^28^. Of the 55,931 sequences that were detectable (87% of the total) at this concentration, 8,412 (15%) bound with *R* ≥ 1.1 (10% enrichment), about half (4,032) of which contained a canonical GCAYG motif. Many oligonucleotides containing the canonical Rbfox binding motifs GCAUG or GCACG were robustly bound, and the distribution of *R* values was shifted upwards with increasing counts of each motif (Figure 1c), consistent with previous work^38^. Similar trends were observed when considering *in vivo* binding, assessed by the density of reads mapping to the genomic region of each oligonucleotide, using published eCLIP data (Figure 1d)^37^. The nsRBNS *R* values of sequences in the library that contained a single GCAUG had a moderate positive correlation with eCLIP read density in the corresponding mRNA regions (*r* = 0.28, P < 1.1 x 10^-25^, Supplementary Figure 1b), as observed previously^38^, confirming that this assay captures some aspects of motif context that are relevant *in vivo*.

### RBFOX2 binds a set of secondary motifs of intermediate affinity *in vitro*

We next aimed to systematically identify sequence elements beyond the “primary” GCAYG motifs that might help to explain the large dynamic range of RBFOX2 binding to the tested oligonucleotides *in vitro*, and the presence of many oligonucleotides with binding above background that lacked GCAYG motifs. We used an iterative enrichment analysis, identifying the most enriched *5*mer (GCAUG), then removing all oligonucleotides containing any instances of this *5*mer and recalculating motif enrichments from the remaining oligonucleotides (Figure 1e). This procedure, similar to that employed in the SKA algorithm for random library RBNS analysis^28^, but adapted to the lower complexity of the naturally-derived sequence library, is designed to prevent the spurious detection of *5*mers that are enriched solely because of overlap with a previously-identified higher-affinity *5*mer. We performed the iterative *R* analysis on data from the highest RBFOX2 concentration, 1.1 µM, which should favor binding to lower-affinity sites^28^.

At each of the first four iterations, the most enriched *5*mers matched the pattern GNNYG. At iterations 5–9, *5*mers matched either GNNUG (GUAUG, GCUUG), NGCAU (UGCAU), or were composed of W (W = A or U) bases only (AAAAA, UUUUU). We considered these as three different classes of potential Rbfox binding motifs, which might bind RBFOX2 directly or might be enriched for other reasons, such as enrichment in the neighborhood of bound motifs in the library. To cast a wide net, we also considered other *5*mers matching these patterns that had *R* values at least 2.5 standard deviations above the mean in the remaining oligonucleotides, adding the GNNUG *5*mers GCCUG and GUGUG, the NGCAU *5*mers CGCAU and AGCAU, and the W_5_ motifs UAUAU and AUAUA to our list of potential binding motifs.

When examined at the oligonucleotide-level, the six GNNUG secondary motifs robustly increase the *R*-value of an oligonucleotide when several motif occurrences are present and while excluding oligonucleotides that contained either primary motif (Figure 1f, P < 7.9 x 10^-29^). The NGCAU and W_5_ *5*mers do so to a somewhat lesser extent (Supplementary Figure 1c-d; P < 8.8 x 10^-28^, P < 0.001, respectively). We also observed increased eCLIP enrichment in regions with increasing numbers of GNNUG motifs, again excluding regions containing either primary motif (Figure 1g, P < 9.6 x 10^-6^)^40^. However, NGCAU and W_5_ *5*mers are not robustly enriched in eCLIP (Supplementary Figure 1e-f, P < 0.071, P < 0.017, respectively). Because of the highly significant binding signal for the six GNNUG *5*mers in both nsRBNS and eCLIP, we chose to pursue only this subclass of secondary motifs for further study.

Together, these six non-GCAYG *5*mers comprised our set of candidate “secondary” Rbfox motifs. These observations, using longer natural sequences containing variable numbers of motifs, support direct recruitment of RBFOX2 to RNAs by these secondary motifs both *in vitro* and *in vivo*.

### Biochemical characterization of Rbfox secondary motifs

We sought to characterize the biochemical features of secondary motifs in nsRBNS. Our iterative method suggested that the G1, G5 and U4 positions of the Rbfox motif are the most critical sites of recognition for the protein. This observation is consistent with published structures of RBFOX1 bound to (U)GCAUG(U), in which the primary hydrogen bonding contacts occur at G1, U4, and G5, while the other bases are recognized mostly by shape (Figure 2a)^41^. The idea that Rbfox tolerates nucleotide variants at positions 2, 3 is also consistent with the conservation pattern of GCAUG in introns and 3’ UTRs, where we observed that the G1 and G5 positions are most strongly conserved, with the intervening positions showing less constraint (Figure 2b). Examining the nsRBNS enrichment of *6*mers ending with the six GNNUG *5*mers, we observed a preference for C or U at the first position, consistent with the preference of GCAYG motifs for an upstream U or C (Supplementary Figure 2a)^41^.

**Figure 2.**
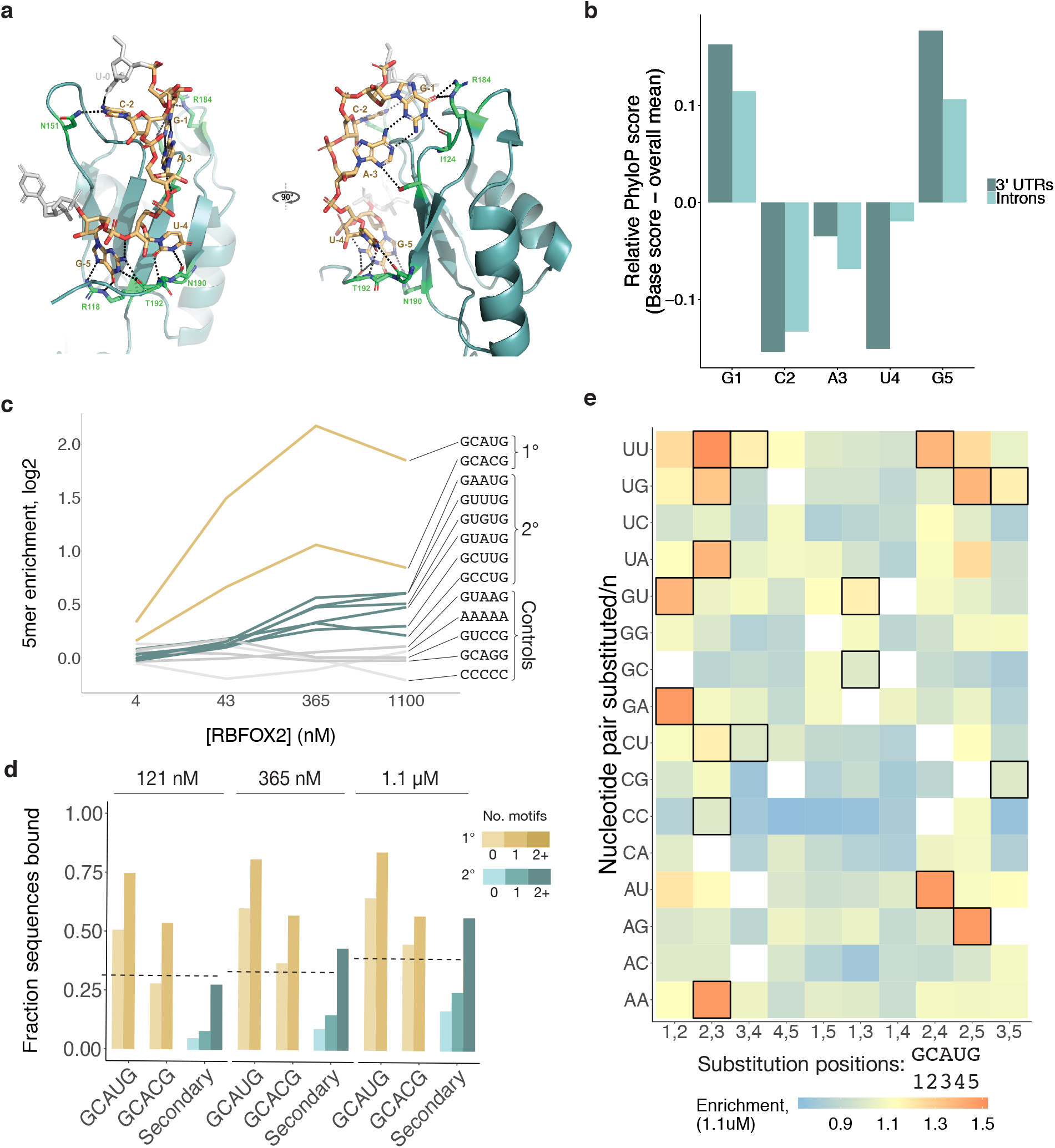
Rbfox proteins reproducibly bind a class of secondary motifs with moderate affinity. a. Ribbon structure of RBFOX1 (teal) bound to UGCAUG (gold). Adapted from Auweter *et al*^41^. Protein–RNA hydrogen bonds are indicated in black. b. Guanines at positions 1 and 5 of the Rbfox primary motif GCAUG are the most strongly conserved bases within the motif. Relative per-base conservation, as represented by PhyloP score, for each position of GCAUG at all instances of the motif in 3’ UTRs (light teal) and introns (dark teal). c. nsRBNS *R* values at 1.1 µM RBFOX2 concentration for all pentamers diverging from GCAUG or GCACG by 1 or 2 bases. Secondary motifs identified here are outlined in black. d. GCAUG and GCACG motifs decline in enrichment at the highest protein concentration. Primary (gold), secondary (teal), polyA and polyC (light grey), and GCAGG, GTAAG, and GTCCG (dark grey) *R* values are shown across four concentrations of RBFOX2 nsRBNS experiments. e. Analysis of fraction of oligonucleotides bound in nsRBNS at three concentrations indicates that six secondary motifs yield binding approximating that of one primary motif at high RBFOX2 concentrations. An oligonucleotide was considered bound if it had an *R* value of at least 1.1.

To further explore these candidate Rbfox motifs, we reanalyzed data from previous random pool RBNS and intronic nsRBNS experiments performed with RBFOX2^28, 38^. In random RBNS, which employs much shorter sequences that are unlikely to contain more than a single secondary motif by chance, secondary motifs were only weakly enriched (Supplementary Figure 2b). However, in an nsRBNS experiment using highly conserved intronic regions, U-rich secondary motifs GCUUG, GUUUG, and GUGUG were enriched (Supplementary Figure 2c). Thus, the length and sequence composition of the library used in nsRBNS may impact ability to detect specific secondary motifs.

Our design with varying concentrations of RBFOX2 enabled analysis of relative RBP occupancies and saturation. Consistent with previous observations^28^, Rbfox primary motifs had high enrichment that declined at the highest protein concentration (1.1 µM), consistent with these sites reaching saturation around the 2^nd^-highest concentration of 365 nM (Figure 2c). Control *5*mers (e.g., AAAAA, CCCCC) or *5*mers differing from GCAUG by one substitution but not identified as secondary motifs (GCAGG, GUAAG, GUCCG) had low and flat *R* values across concentrations, consistent with absence of specific binding. In contrast, the six secondary motifs identified above exhibited moderate but steadily increasing enrichment at successive protein concentrations, flattening somewhat at the highest level, consistent with moderate binding affinity that becomes saturated only at or beyond 1.1 µM [RBFOX2]. We noted that about half of the oligonucleotides containing six or more secondary motifs but no primary motifs were bound at 1.1 µM [RBFOX2], comparable to the fraction of oligonucleotides containing a single GCAUG that were bound at lower Rbfox concentrations (Figure 2d). This observation suggests that clusters of secondary motifs (which turn out to be common in the transcriptome) may function in regulation similarly to single primary motifs if Rbfox concentrations are high. Notably, in a previous study using surface plasmon resonance, several secondary motifs had K_d_ values in the 100-600 nM range^29^, well above the low nanomolar value observed for GCAUG, but comparable to the highest affinity motifs of many other RNA-binding proteins^42^. Of the 4,358 sequences bound at *R* ≥ 1.1 in nsRBNS that lacked a primary motif, 1,885 (43%) contain at least one of the ten GNNUG secondary motifs. Thus, the six secondary motifs identified above appear to explain a plurality of the non-canonical (non-GCAYG) RBFOX2 binding to our nsRBNS library.

To ask whether other related *5*mers beyond those identified above could bind RBFOX2, we examined the binding at 1.1 µM [RBFOX2] of all *5*mers with one or two positions substituted from the primary binding motif GCAUG (Figure 2e). The most enriched *5*mers mostly consisted of the two primary and six identified secondary motifs. Besides these six motifs, certain AU-rich *5*mers with two deviations from GCAUG (UUAUG, AUAUG, GUAUU, and GUAUA) also showed modest enrichment.

### RBFOX2 binds to specific secondary motifs *in vivo*

We next asked whether Rbfox binding to individual secondary motifs could also be observed *in vivo*. We analyzed published high-resolution *in vivo* binding data from mouse embryonic stem cells (mESCs) generated using the individual-nucleotide resolution CLIP (iCLIP) assay^27^. We constructed a meta-motif plot summing read density as a function of distance for every instance of a given *5*mer in expressed introns or 3’ UTRs (Figure 3a). Primary motifs GCAUG and GCACG, and all six GNNUG secondary motifs showed a characteristic peak located somewhat 3’ of the motif location, as expected^28^, with broader peaks observed in a few cases (e.g., GAAUG). Some of the motifs also showed evidence of crosslinking at U4 of the motif (Supplementary Figure 3a). However, the four AU-rich *5*mers flagged above in the high [Rbfox] nsRBNS data lacked similarly robust peaks and thus discarded in downstream analyses (Supplementary Figure 3b). In general, the peaks were stronger and more clearly defined in introns than in 3’ UTRs, likely reflecting the predominantly nuclear localization of RBFOX2 and its prominent role in splicing. We generated a CLIP enrichment value for each motif by normalizing the read density at the peak apex by the read density at positions distal from the peak (Supplementary Figure 3c). Comparing 3’ UTR iCLIP *5*mer enrichments with nsRBNS *5*mer *R* values in introns (Figure 3b) and 3’ UTRs (Supplementary Figure 3d) yielded similar results: GCAUG and GCAUG-overlapping *5*mers were the most highly enriched by both measures, with GCACG close behind, and the six secondary motifs near the top of remaining *5*mers by both measures.

**Figure 3.**
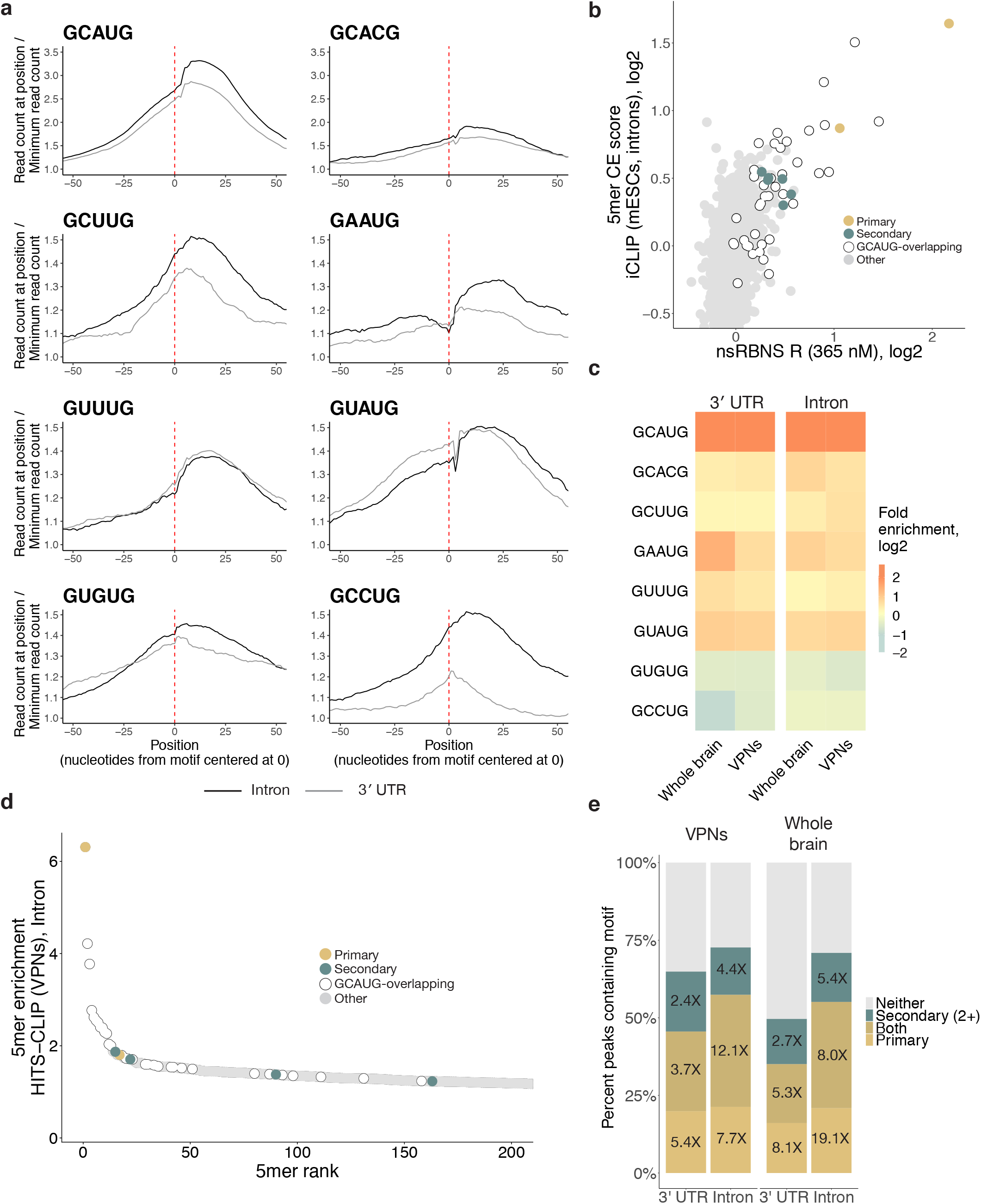
Rbfox proteins bind secondary motifs *in vivo*. a. Both primary and secondary motifs show characteristic read peaks near 0 in a metaplot centered at the motif in introns (black) and 3’ UTRs (grey) in RBFOX2 iCLIP data^27^. iCLIP reads containing the motif of interest were aligned with position one of the pentamer at 0 and normalized to the minimum read count in an 80-nt window (50-nt window shown). Y-axis range was reduced for secondary motifs. b. Correlation of intronic iCLIP- and nsRBNS-enriched *5*mers. CLIP enrichment (CE) scores were computed for iCLIP peaks by dividing the read count in the highest five base window by the read count in the lowest five base window. Secondary motifs indicated in teal, primary motifs indicated in gold. Grey dots indicate “hitchhiking” motifs that overlap the primary motif GCAUG by at least three bases but do not have intrinsic Rbfox affinity. c. RBFOX1 HiTS-CLIP data^8, 14^ shows enrichment for primary motifs GCAUG and GCACG and secondary motifs GCUUG, GAAUG, GUUUG, and GUAUG in both 3’ UTRs (top) and introns (bottom) as compared to transcriptomic frequencies, while secondary motifs GUGUG and GCCUG appear depleted (blue). Enrichment was calculated based on the *5*mer composition of 100-base CLIP peak regions centered around the apex of the CLIP peak relative to *5*mer composition of transcriptomic *5*mer content. d. Four secondary motifs (GCUUG, GAAUG, GUUUG, GUAUG) are among the top 200 highly enriched *5*mers derived from intronic HiTS-CLIP peaks from mouse ventral spinal neurons. Primary motifs in gold, secondary motifs in teal. Peaks calculated as above. e. High-confidence CLIP peaks in two different Rbfox1 HiTS-CLIP datasets attributable to primary (light gold), or secondary (teal), or both (dark gold) motifs in both 3’ UTRs and introns. 100-base CLIP peak regions centered around the apex of the CLIP peak were searched for presence of a primary or two or more secondary motifs in that region. Fold enrichments above transcriptomic background are indicated by text in the corresponding bars. Peaks containing neither primary nor two or more secondary motifs are shown in grey.

We sought to further explore *in vivo* binding to specific *5*mers in neuronal cell types, where expression of Rbfox protein is often very high. Using two high-throughput crosslinking and immunoprecipitation (HiTS-CLIP) datasets – in whole mouse brain and *in vitro*-differentiated ventral spinal neurons (VPNs)^8, 14^ – we analyzed the enrichment of secondary motifs in stringently filtered CLIP peaks present in two replicates, examining both 3’ UTR and intronic regions. As expected, primary motifs GCAUG and GCACG were strongly (4.1- to 6.2-fold) and moderately (1.3- to 1.7-fold) enriched, respectively, and GCAUG was the most enriched *5*mer in each HiTS-CLIP dataset (Figure 3c-d, Supplementary Figure 4). Notably, four of the six secondary motifs (GCUUG, GAAUG, GUUUG, GUAUG) were also enriched in CLIP peaks to extents approaching that observed for GCACG, indicating association with Rbfox protein *in vivo*. The magnitude of enrichment in 3’ UTRs and introns was similar for most motifs. Because secondary motifs GUGUG and GCCUG lacked robust enrichment in HiTS-CLIP data, we chose to exclude them from further analysis of *in vivo* function, focusing instead on the remaining four motifs GCUUG, GAAUG, GUUUG, and GUAUG.

We explored the extent to which secondary motifs can account for CLIP peaks not explained by the presence of a primary motif. To do this, we looked for instances of primary, secondary, or both motif types within 50 bases upstream and downstream of the apex of HiTS-CLIP peaks that appeared in both replicates of 3’ UTR or intronic HiTS-CLIP data, analyzing two different datasets (Figure 3e)^8, 14^. In both datasets, ∼50% of peaks contained a primary motif or both types of motifs (as seen previously^23^), while another 15-20% of peaks lacked primary motifs but contained two or more secondary motifs. Even in the absence of primary motifs, presence of at least two secondary motifs was enriched approximately five-fold in the intronic HiTS-CLIP peaks, and about 2.5-fold in 3’ UTR peaks. A minimum of two secondary motifs was required in this analysis to compensate for the higher frequency of these motifs in the transcriptome. Thus, consideration of secondary motifs helps to explain a substantial additional fraction of CLIP peaks that lack a primary motif.

### Secondary motifs regulate splicing in an Rbfox-dependent manner

We next sought to assess the potential of secondary motifs to mediate Rbfox-dependent regulation. We adapted a bichromatic splicing reporter (pRG6)^43^ by introducing one copy of GCAUG or six copies of one of three secondary motifs (GCUUGx6, GAAUGx6, or GUUUGx6) into a 250 nt region downstream of an alternative exon (Figure 4a). For the secondary motifs, the six copies were inserted into six interspersed positions (see Methods), and a control vector was constructed for each by inserting a permuted version of the motif at the same positions (designated as pGCUUG, etc., with “p” for permuted). In this reporter design, exclusion of the alternative cassette exon yields a transcript in which the dsRED ORF is in frame, while inclusion of the exon shifts the frame so that EGFP is produced. Thus, the effects of different Rbfox motifs on exon inclusion can be assessed by measuring red versus green fluorescence. We co-transfected the modified RG6 plasmid with or without a Cerulean:RBFOX1 fusion protein (pCERU:rbFOX1) into HEK293T cells to augment low endogenous levels of RBFOX2 protein.

**Figure 4.**
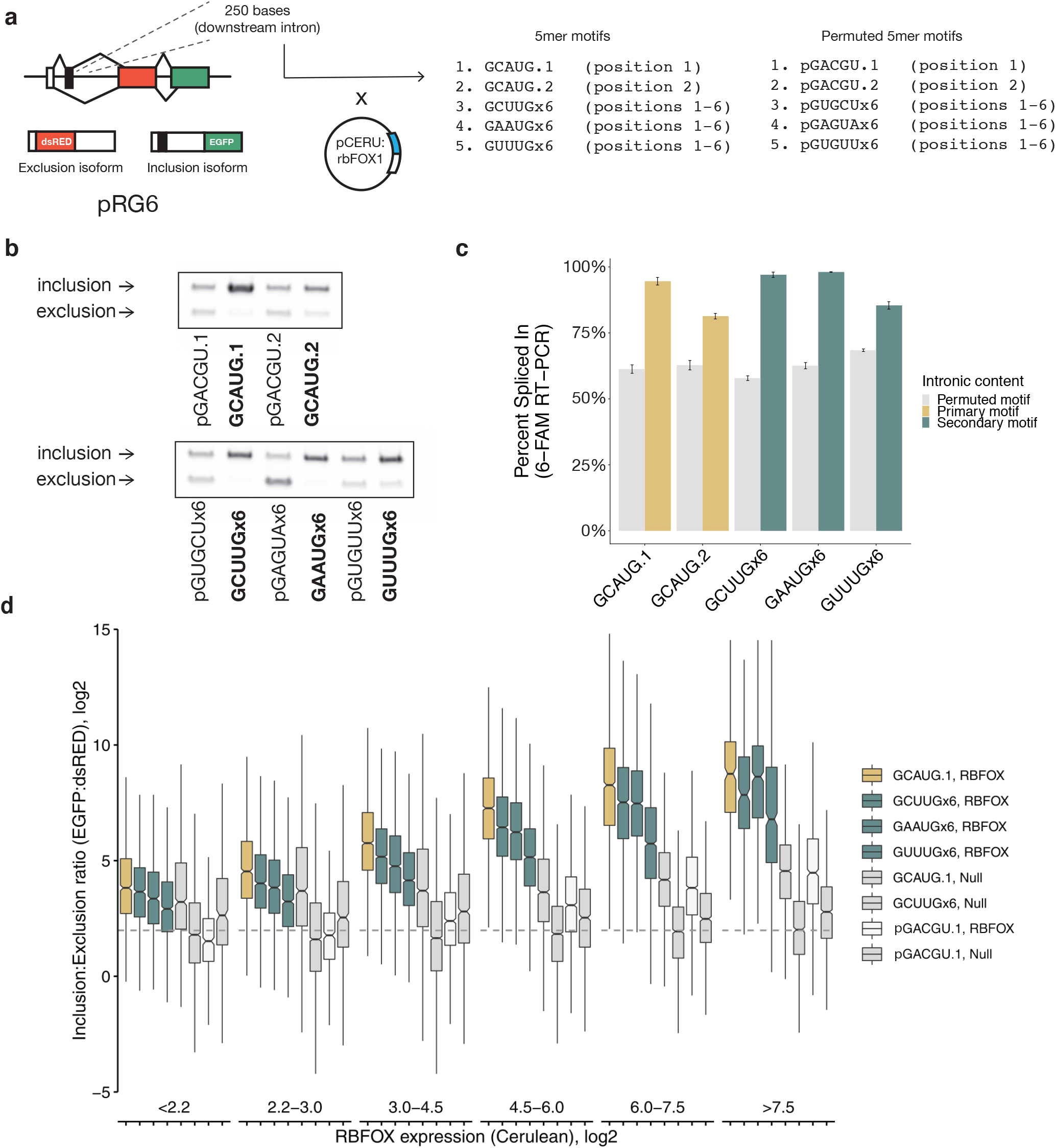
Secondary motifs in downstream introns promote exon inclusion in an Rbfox-dependent manner in a splicing reporter. a. Experimental design of *Rbfox1* splicing reporter. One GCAUG primary motif or six copies of a secondary motif were cloned in a 250-base window downstream of an alternative exon in the RG6 dual fluorescent splicing reporter. Plasmids were co-transfected in HEK293T cells with a plasmid expressing fluorescently labelled RBFOX1 to monitor cellular protein levels. b. Semi-quantitative PCR with 5’ 6-FAM-labelled primer shows robust exon inclusion in the presence of both primary and secondary motifs in an *Rbfox1*-dependent manner. c. Percent spliced in (PSI) values of primary motifs (gold), secondary motifs (teal), or motif permutations (grey) in the downstream intron after overexpression of *Rbfox1*. Error bars show SD of triplicate experiments. For GAAUG, the median permutation value was used due to the introduction of a splicing silencer in its permuted form. d. Six secondary motifs approximate the activity of one primary motif at higher RBFOX1 concentrations. The indicated RG6 plasmids, including scrambled motif controls (light grey), were co-transfected with fluorescently labelled RBFOX1 (or CFP alone, medium grey) into HEK293T cells and monitored by flow cytometry to assess the inclusion isoform (GFP), exclusion isoform (dsRED), and RBFOX1 (Cerulean) expression of single cells.

To measure the effect of Rbfox-mediated regulation of splicing in the presence of secondary motifs, we measured exon inclusion by RT-PCR with a fluorescently-labeled primer (Figure 4b). Insertion of either a single GCAUG or six copies of any of the secondary motifs tested drove percent spliced in (PSI) values of the exon to nearly 100% in the presence of exogenous RBFOX1 (Figure 4c). Notably, presence of a single GCAUG was sufficient to drive high inclusion without exogenous Rbfox, while presence of secondary motifs exerted strong effects on splicing predominantly when exogenous Rbfox protein was added via co-transfection with the pCERU:rbFOX1 plasmid (Supplementary Figure 5a). This observation is consistent with the expectation that higher Rbfox levels are required for effective binding of secondary motifs.

We observed similar results for protein isoform expression across single cells. Using Cerulean fluorescence to quantify (exogenous) RBFOX1 levels, and the EGFP:dsRED ratio to quantify exon inclusion, we were able to directly measure the relationship between RBFOX1 levels and exon inclusion driven by secondary motifs (Figure 4d, Supplementary Figure 6). Generally speaking, six copies of any of the secondary motifs GCUUG, GAAUG, or GUUUG drove exon inclusion to about the same extent as a single copy of the primary motif GCAUG. At low levels of RBFOX1 expression (bins 1-2), all constructs exhibited exon inclusion similar to the GCAUG construct co-transfected with an empty Cerulean vector (GCAUG.1, Null), confirming the RT-PCR results. As RBFOX1 expression increases, constructs containing primary or secondary motifs increased exon inclusion well above background. Exon inclusion did not increase or increased modestly in control constructs (Methods). Across two replicates, inclusion was significantly more correlated with Rbfox expression when Rbfox primary or secondary motifs were present (Supplementary Figure 5b). Together, these observations demonstrate that secondary motifs can mediate robust regulation of splicing, particularly when levels of Rbfox proteins are high.

### Secondary motifs are associated with splicing regulation at specific stages of neuronal differentiation

Given our observations that secondary motifs are most effectively bound by Rbfox in regimes of high Rbfox expression, we examined neuronal differentiation, a system in which high expression levels of Rbfox occur naturally. In an eight-point time course of mESCs differentiating into glutamatergic neurons^39^, Rbfox increases five-fold in expression from the radial glia stage (RG) to developmental stage 3 (DS3) (Figure 5a). In this time course, cells progress from a bipolar-shaped progenitor neuron (RG) to fate-specified developmental stage 1 (DS1) neurons, then to developmental stage 3 (DS3) over a seven-day period, and eventually become mature neurons after substantial growing and pruning of dendrites (Figure 5a). We examined the relationship between Rbfox-induced exon inclusion with the presence of primary or secondary motifs in the first 250 nt of the downstream intron. We first looked at the interval RG– DS1, where Rbfox increases to moderate expression, and DS1–DS3, where Rbfox reaches its highest levels in the time course. For primary motifs GCAUG and GCACG, exon inclusion is significantly correlated with motif frequency at both intervals. However, the secondary motifs are correlated with exon inclusion only at the DS1–DS3 interval, suggesting that they are mostly active later, when Rbfox levels are higher (Figure 5b).

**Figure 5.**
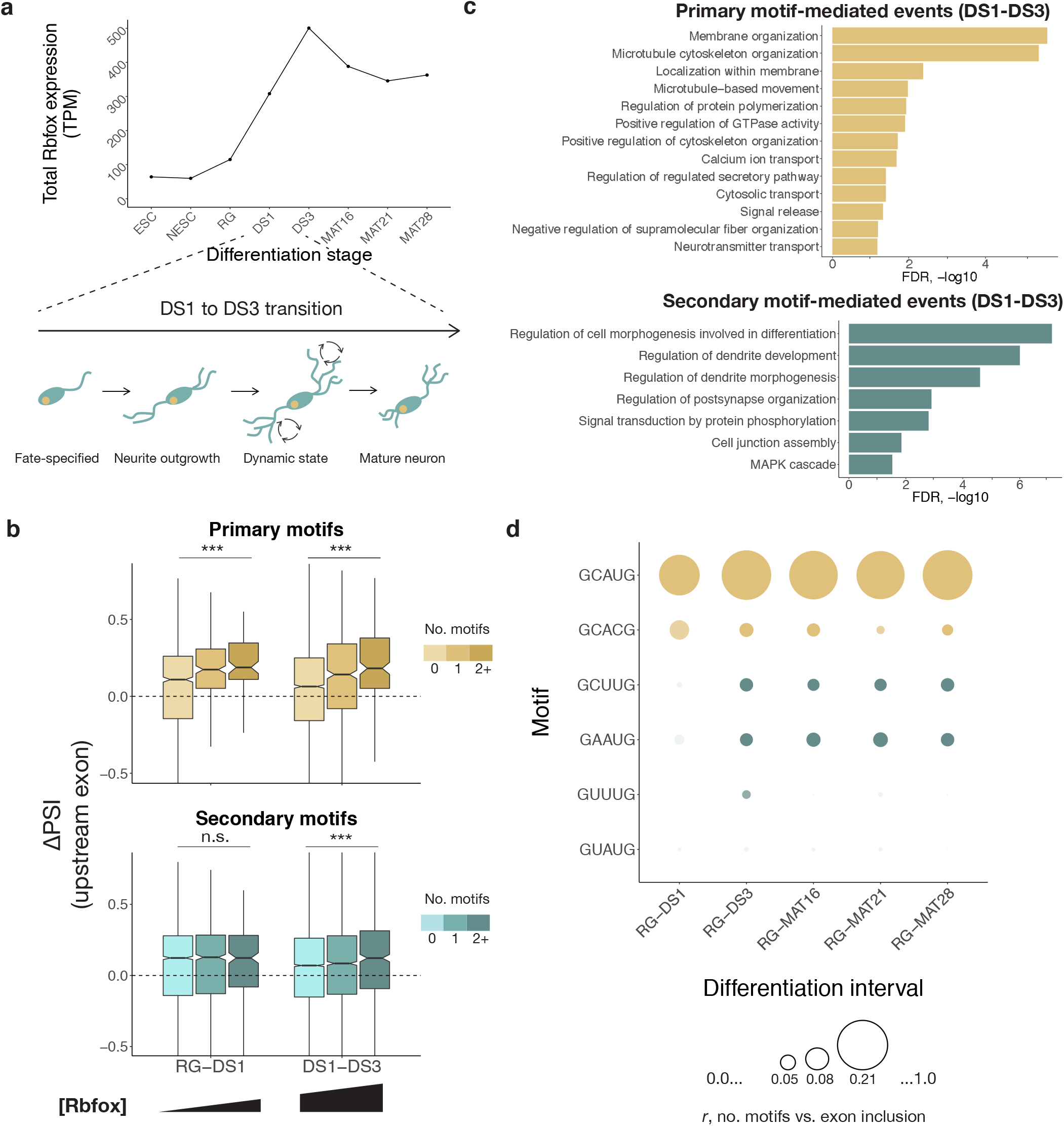
Secondary motifs enable splicing regulation at distinct concentration Rbfox concentration ranges in neuronal differentiation. a. Total expression of *Rbfox1*, *Rbfox2*, and *Rbfox3* in transcripts per million (tpm) based on RNA-seq during a neuronal differentiation time course^39^. b. Count of Rbfox primary motifs (gold shades) in the downstream intron is correlated with delta PSI at both low-moderate (RG-DS1) and moderate-high (DS1-DS3) transitions of Rbfox expression during *in vitro* neuronal differentiation, while Rbfox secondary motif count (teal shades) is correlated with Rbfox activity only at the moderate-high Rbfox expression transition, DS1-DS3. Increased color intensity represents 0, 1, or 2+ motifs in the downstream intron. c. Splicing events driven by primary motifs differ in Gene Ontology categories from those driven by secondary motifs within neuronal differentiation. Events were compared to all expressed genes at DS3 and filtered for FDR < 0.1, B > 99, and b > 9. d. Secondary motifs remain engaged throughout subsequent stages of neuronal differentiation. Pearson correlation of secondary motif presence with delta PSI is shown at intervals of neuronal differentiation from radial glia stage to mature 28-day glutamatergic neurons. Size of point indicates correlation coefficient, intensity indicates p-value < 0.05.

In order to examine potential functions of splicing changes in the DS1-DS3 interval, we assessed the functions of genes containing skipped exons whose splicing changed during this interval. For exons associated with primary Rbfox motifs (*n*=388), Gene Ontology analysis yielded enrichment for functions related to membrane and cytoskeletal organization, while exons associated with secondary motifs (*n*=561) gave enrichment for functions in dendrite development and signal transduction (Figure 5c). This distinction likely reflects a regulatory program in which primary motifs mediate earlier splicing events related to neurite outgrowth and secondary motifs mediate a later wave of splicing changes related to dendrite development and signaling, as Rbfox protein levels increase.

To further assess the specific intervals at which splicing regulation occurred, we examined the correlation between motif count and difference in exon inclusion (delta PSI) across every interval of the time course (Supplementary Figure 7). The DS1–DS3 interval was the only interval in which secondary motifs were significantly correlated with Rbfox activity. This interval may represent a distinct transition in the usage of secondary motifs in neuronal differentiation concordant as Rbfox expression peaks. Examining intervals from the RG stage to subsequent stages of the time course (Figure 5d), secondary motifs GCUUG and GAAUG had signal throughout the rest of the differentiation, while GUUUG had signal only at DS3, and GUAUG lacked detectable signal in this analysis. These observations suggest that GCUUG and GAAUG are the most active of the secondary motifs in this system. Other families of RBPs with related binding motifs are also active during neuronal development. Proteins of the CELF family could potentially bind GU-rich motifs such as GUUUG, and Musclebind-like (Mbnl) family proteins could bind to secondary motifs containing GCU or GCC^26^, potentially modulating these correlations.

### Rbfox expression-dependent splicing via secondary motifs in differentiated neuronal cell types

We next asked to what extent Rbfox might contribute to diversification between differentiated neuronal cell types. We analyzed data from thirteen cell types and sub-types that span almost three orders of magnitude of Rbfox gene expression, combining mRNA levels of *Rbfox1*, *Rbfox2* and *Rbfox3*^4^. Specifically, we compared mean differences in PSI between cells grouped by their Rbfox expression from lowest (EC, TRCbitter, OSN25wk) to highest cell types (CGN) (Figure 6a). We expected that the number of GCAUG motifs downstream of alternatively spliced exons should be generally correlated with exon inclusion, whereas secondary motifs should contribute only in cell types with higher Rbfox expression. Indeed, we observed that downstream GCAUG motif count is significantly correlated with exon inclusion, even when comparing medium to low Rbfox expression (P < 0.0015), and is more strongly correlated when comparing high to medium (P < 1.3 x 10^-19^) and highest to low (P < 2.4 x 10^-23^), supporting that our grouping of cell types captures Rbfox concentration-dependent regulation of splicing (Figure 6b). Consistent with our previous results, the strongest secondary motifs GCUUG, GAAUG, GUUUG, and GUAUG together contribute significantly to increased exon inclusion in high (P < 7.7 x 10^-4^) and highest (P < 1.3 x 10^-4^) Rbfox-expressing cells compared to low-expressing cells. When comparing highest to low Rbfox-expressing cells, the slope of the regression predicts that, on average, each downstream GCAUG increases exon PSI by 15%, while each secondary motif yields a 3.6% increase in PSI. Thus, in endogenous loci as in our reporter assays, a small group of secondary motifs is sufficient to confer the regulatory activity of a canonical primary motif.

**Figure 6.**
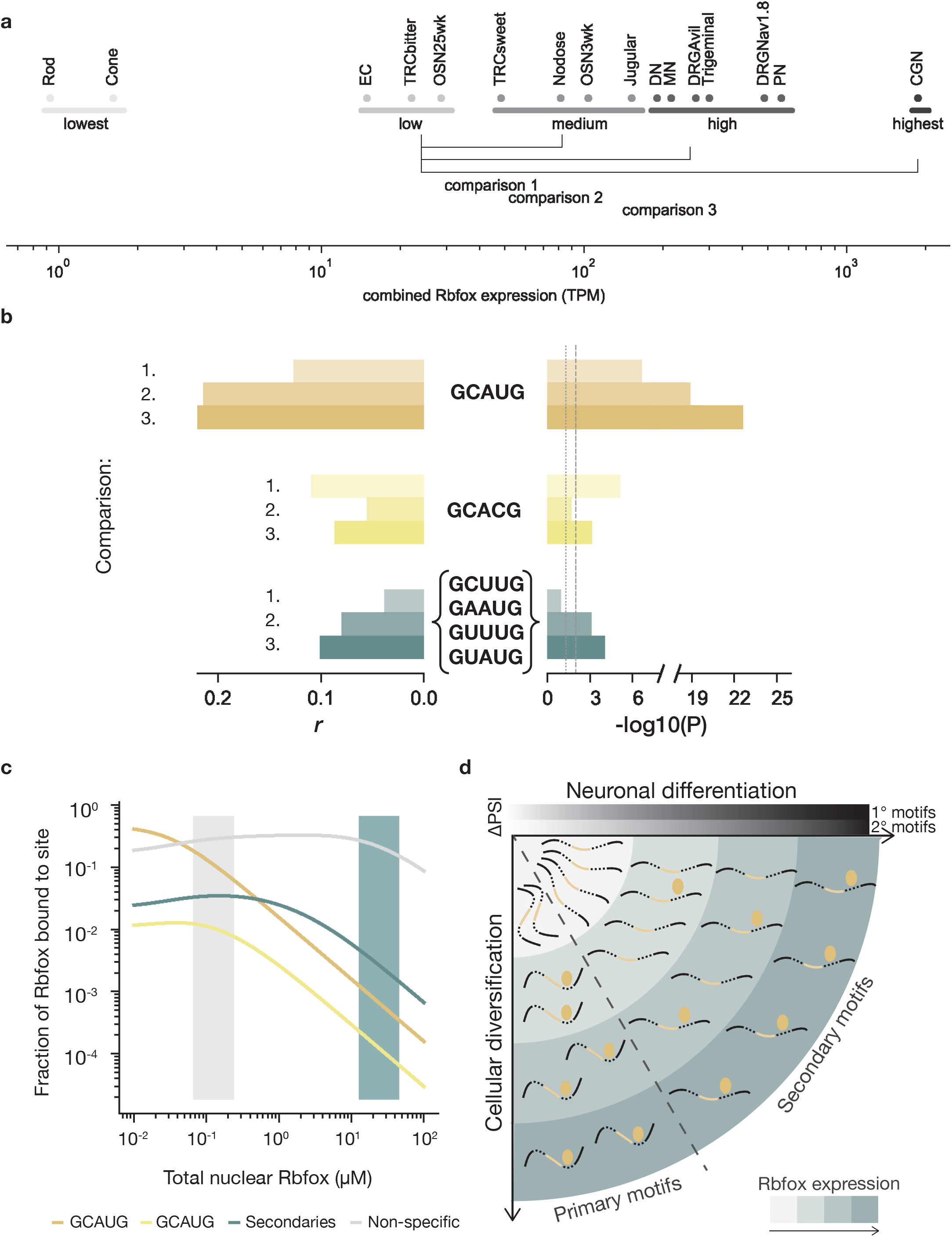
Secondary motifs are active in neuronal cell types with high Rbfox expression. a. Differentiated neuronal cell types from Weyn-Vanhentenryck *et al*.^4^ arranged by combined Rbfox1, Rbfox2, and Rbfox3 expression (log10 of sum of RPKM values). Cell types were grouped into Lowest, Low, Medium, High, and Highest Rbfox expression categories. Cells analyzed: olfactory sensory neurons (OMP+) (OSN), enterochromaffin cells (EC), taste receptor cells (TRC), rod or cone photoreceptors, jugular or nodose visceral sensory ganglia, dopaminergic neurons (DN), motor neurons (MN), dorsal root ganglia sensory neurons (Nav1.8+ or Avil+) (DRG), trigeminal ganglia, Purkinje neurons (PN), and cerebellar granule neurons (CGN). b. Linear regression was performed on 1,909 alternatively spliced exons comparing the Rbfox expression groups indicated in a. Horizontal bars represent the *r* value (left) and significance (right) of the correlation between the number of occurrences of different groups of motifs (middle) and ΔPSI values between Medium and Low (1.), between High and Medium (2.), or between Highest and Medium (3.). Rbfox expression regimes. c. An equilibrium model for Rbfox binding to intronic sequences in the nucleus predicts that a substantial fraction of protein is associated with secondary motifs (teal), at high expression levels of Rbfox (teal shaded area), whereas the primary motif GCAUG (gold) dominates specific binding at low to moderate expression levels of Rbfox (grey shaded area), but is limited by saturation of these sites at high levels. GCACG (yellow, variant primary motif) is too rare to capture a large fraction of Rbfox. Non-specific *5*mers (grey) may capture a substantial fraction of Rbfox due to their collectively very high abundance. d. Graphical summary of how Rbfox proteins (golden ellipses) regulate distinct splicing events at different expression levels (teal shading). Secondary motifs become active only at higher Rbfox levels, occurring at later stages of neuronal differentiation or in high-Rbfox cell types, while primary motifs become active at earlier stages and in cell types with medium as well as high Rbfox levels.

### A quantitative model for Rbfox expression-dependent, differential motif activity

In order to better understand the distribution of Rbfox protein binding across sites in the nuclear transcriptome of a mammalian cell, we built a quantitative equilibrium model (Supplementary Figures 9-10). We used standard estimates of the ranges of cell and nuclear volume, and of mRNA abundance (10^5^ to 10^6^ molecules per cell) and half-life (several hours), estimated the rate of mRNA production required to balance decay, and used this value to estimate the abundance of intronic RNA in the nucleus, assuming rapid turnover of introns (half-life of 1 min). To illustrate the plausible range of RNA concentration in the nucleus, we consider two scenarios at opposite ends of this range: a large cell with lower RNA turnover and RNA concentration (Figure 6c, Supplementary Figure 10a,b) and a small cell with higher RNA turnover and RNA concentration (Supplementary Figure 10c,d) (Methods).

Using gene expression data from the DS3 differentiation stage with available genome annotations and the model above, we estimated the aggregate concentration intronic RNA *5*mers as between ∼14 and 56 μM (Methods). Of this total, the concentration of GCAUG will fall between ∼16 to 63 nM, with GCACG fivefold lower (3-12 nM), and the four secondary motifs together occurring at severalfold higher concentration (67 to 270 nM) (Supplementary Figure 10a). After assigning approximate dissociation constants to all Rbfox *5*mer motif variants by calibrating our random RBNS data for human RBFOX2 and RBFOX3^26^ to SPR data^29^ (Supplementary Figure 9, Supplementary Table 3), we modeled the equilibrium distribution of Rbfox proteins across the pool of nuclear binding sites, analogous to a previous model for miRNAs^44^. We show our predictions as a function of nuclear Rbfox concentration (Figure 6c). Next, we estimated the nuclear Rbfox concentration from the range of Rbfox mRNA expression across neuronal cells (Figure 6a), for a proteins-per-mRNA ratio of 2,800 – 10,000, in line with a reported protein:mRNA ratio for *Rbfox2.*^45^ In cells with low to intermediate Rbfox expression, between 20% to 40% of Rbfox protein is recruited to primary motifs even though they represent a small fraction of the available RNA pool. On the other hand, at very high Rbfox expression, the primary motifs become saturated (due to their high affinity and low abundance) and a larger fraction of Rbfox protein occupies secondary motifs (Figure 6c, Supplementary Figure 10d). We estimate that individual instances of secondary motifs reach ∼50% occupancy at nuclear Rbfox concentrations between 10 and 50 μΜ (Supplementary Figure 10b,c). We note that our estimates of affinity for non-specific (background) *5*mers from random RBNS data likely represent lower bounds on the K_d_ and that a higher non-specific K_d_ would yield both more free Rbfox and greater occupancy of secondary motifs. Using different gene expression profiles (RG, DS1, DS3) had a marginal impact on these findings, presumably because the average intronic composition does not vary much between neuronal cell types and conditions (data not shown).

We conclude that the secondary motifs identified here contribute to regulation under physiological conditions with greatest activity at high nuclear concentrations of Rbfox (Figure 6d).

## Discussion

The Rbfox family of splicing factors play important roles in development and maintenance of various tissues and are among the best-studied RBPs. Even so, aspects of their RNA binding and regulatory activity are incompletely understood. Here, we have shown that Rbfox family proteins bind a defined set of abundant secondary motifs *in vitro* and *in vivo*. These motifs enable concentration-dependent regulation of exon inclusion by Rbfox in neuronal differentiation and cellular diversification, adding a second wave of regulation in cells that achieve high Rbfox expression. The mediation of temporally-staged and cell type-specific activity by use of secondary motifs has rarely been observed for RBPs^46, 47^, but may be reasonably common.

Though variant motifs have occasionally been noted in other studies of Rbfox proteins, many studies go on to filter out non-GCAUG elements in their datasets^8–10, 12– 14, 34^. In some cases, binding of secondary motifs may have been attributed to other proteins, e.g., presence of GUGUG in Rbfox CLIP peaks has been attributed to binding by SUP-12^32^ or HNRNP M of the LASR complex^33^. Certainly, the potential for direct binding of GUGUG motifs by Rbfox proteins should also be considered. Additionally, our findings that secondary motifs can contribute to neuronal diversification may explain varying enrichments of primary and secondary motifs among CLIP data derived from different cell lines^8, 14, 33^.

Suboptimal motifs may function in a few different ways. First, they may enhance binding to nearby primary motifs. Second, a cluster of several secondary motifs may create a stretch of sequence with total Rbfox binding comparable to that of a high-affinity motif. Consistent with this idea, we observed increased binding to oligonucleotides with larger numbers of secondary motifs in nsRBNS, and found evidence supporting that 4-6 copies of a secondary motif can enhance exon inclusion to an extent comparable to that of a single GCAUG motif in vivo. More generally, suboptimal motifs can function as *cis-*regulatory elements that function only at high levels of a *trans-*factor to narrow the temporal or spatial scope of activity, an effect we observed for Rbfox regulation in cellular differentiation and diversification.

Examples of spatial and temporal regulation via secondary motifs have been described for several transcription factors (TFs). The *C. elegans* TF PHA-4 activates its primary motif target in early pharyngeal development, while its secondary targets are activated only later, when protein levels are higher^49^. In mice, the TF PREP1 activates its enhancers sequentially according to their binding site affinity during eye lens development^50^, and suboptimal motifs for the GATA and FGF families of TFs enable tissue-specific expression in tunicates^48^. Recent work has demonstrated that secondary motifs can be activated when levels of a TF are boosted by gene amplification, e.g., the MYC oncoprotein binds low-affinity motifs in a sequence-specific and dose-dependent manner in tumors in which the *Myc* locus is amplified, which may contribute to transcriptional misregulation^51^. At the genomic level, Bayesian biophysical models that considered binding to both high- and low-affinity sites better predicted gene expression in human cells, supporting the importance of low-affinity sites to regulation^52^.

Our study argues that it is time to reconsider the simplifying dichotomy of “binding site” versus “non-binding site” and to start to consider site affinities, extending consideration to sites of lower affinity when RBP activity is high^53^. For example, our models indicate that relatively low abundance primary motifs may become saturated at physiological protein concentrations. Proteins with high affinity and specificity like Rbfox proteins (and also Pumilio, Muscleblind, Musashi, and certain others) – especially those expressed at variable and high levels – may be fundamentally different from RBPs (such as SR proteins) that bind fairly degenerate primary motifs that occur in quantities sufficient to titrate levels of the RBP. The FACS-based approach introduced here may be useful for studying the relationship between RBP concentration and splicing regulation.

Taken together, our experiments and computational analyses demonstrate that secondary Rbfox motifs contribute significantly to Rbfox-dependent gene regulation, with their relatively low affinity enabling highly cell-type and stage-specific regulation. This behavior, observed at high overall levels of Rbfox, could also become relevant in regimes of high local Rbfox concentration. For example, Rbfox has been associated with phase-separated compartments in the nucleus^22^, an additional context where high concentration may drive binding of secondary motifs.

## Experimental procedures

### Cloning, expression, and purification of RBFOX2

The RRM domain of *RBFOX2* (amino acids 100-194) was cloned into the pGEX6P-1 expression vector (GE Healthcare, 28-9546-48) downstream of a GST-SBP tandem affinity tag. Following 12 h recombinant expression in Rosetta Competent Cells (Millipore, #70954) at 12° C with ampicillin and chloramphenicol selection, the protein was expressed by addition of IPTG and purified via the GST tag as described previously^26, 28, 38^.

### Library design of natural 3’ UTR sequences

GENCODE^54^-annotated human transcripts were evaluated for presence of appropriate start codon and stop codon sequences at the corresponding GENCODE-annotated positions. Coding genes that had a 3’ UTR between 100 and 10,000 bp were considered for library inclusion. For the *H. sapiens* library, 120 transcript pairs were included because of evidence of alternative 3’ UTRs, 720 transcripts were selected based on expression level of >10 fragments per kilobase million (FPKM) in both HepG2 and K562 cell lines, and 360 other transcripts were selected at random. Each of the resulting transcripts was assigned a homologous *M. musculus* transcript by identifying a transcript in the homologous *M. musculus* gene in which the annotated polyA tail was ±150 nt from the location of the homologous *H. sapiens* polyA tail site, as identified by Batch Coordinate Conversion (liftOver, UCSC Genome). This procedure generated 1108 *H. sapiens* 3’ UTRs paired with 1104 *M. musculus* 3’ UTRs. Each 3’ UTR sequence was then split into overlapping 110 nt segments at 43 nt intervals to achieve approximately 2.5X coverage of each 3’ UTR, yielding 64,319 unique sequences.

### Natural sequence RNA Bind-n-Seq procedure and analysis

The 64,319-oligonucleotide library was synthesized by Twist Biosciences and nsRBNS was performed as previously described^26, 28, 38^. Briefly: Library was amplified with Phusion Polymerase (NEB, #E0553L, primers below), *in vitro*-transcribed, treated with Turbo DNase (Thermofisher, #AM2238), and gel- and phenol:chloroform:isoamyl alcohol-purified. Streptavidin T1 magnetic beads (Invitrogen, #65601) and 250 nM of the prepared library was incubated with recombinant, tagged RBFOX2 at concentrations of 0 (no protein control), 4, 14, 43, 121 (2 replicates), 365, and 1100 nM for 1 hr at 4° C to equilibrium binding in binding buffer (25 mM Tris, 150 mM KCl, 0.1% Tween, 0.5 mg/ml BSA, 3 mM MgCl_2_, 1 mM DTT, pH 7.5). RBP:RNA:bead complexes were pulled down with a magnet and washed gently twice with wash buffer (25 mM Tris, 150 mM KCl, 0.1% Tween, 0.5 mM EDTA, pH 7.5). RNA was eluted by two separate incubations with 4 mM biotin (pH 7.5) for 30 min at room temperature. RNA eluate was purified with Ampure beads (Beckman Coulter, #A63987), and the resulting RNA was reverse transcribed with Superscript III (Thermofisher, #18080093, primer below). The cDNA was amplified for 6-16 cycles (primers below) with Phusion Polymerase (NEB, #E0553L) and gel-purified with ZymoClean Gel DNA Recovery Kit (Genesee Scientific, #11-300C) to produce the final library. Each library was sequenced single-end on an Illumina HiSeq 2500 instrument. Sequences with at least 100 associated reads in the input sample were considered for further analysis. Read counts in each sample were normalized by the total reads in that sample. To produce enrichment (*R*) values at the single-oligonucleotide level, normalized reads for an oligonucleotide in a given sample were divided by the normalized reads for that oligonucleotide in the 0 nM control sample. In general, sequences enriched at an *R* value of 1.1 (10% enrichment) were considered “bound”.

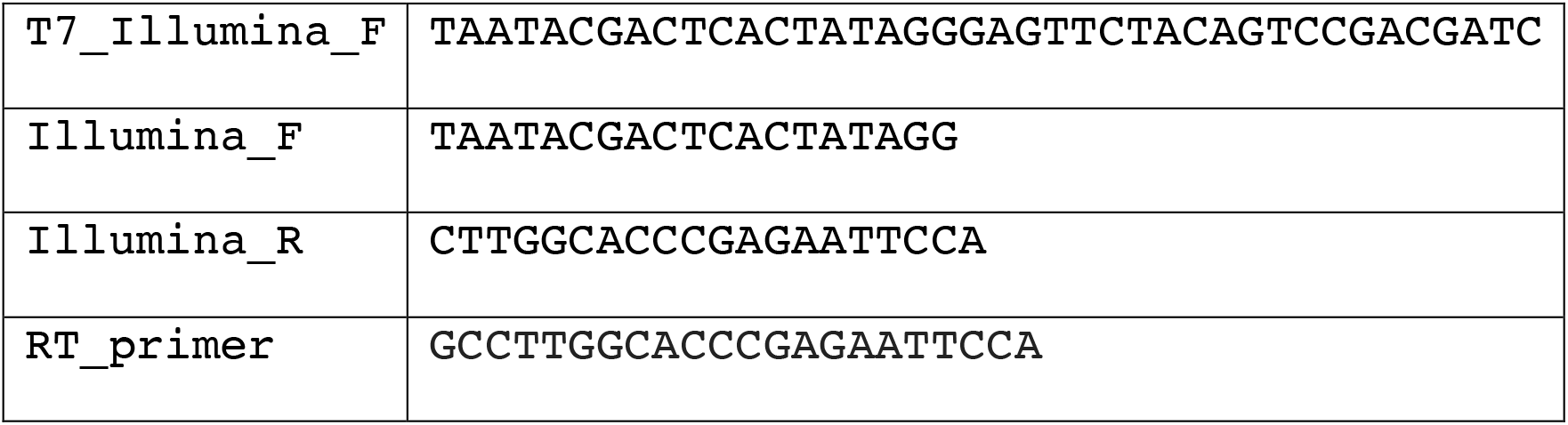

### PhyloP scores

Plus-strand hg19 46-way alignment phyloP^55^ scores were obtained for each genomic position from the UCSC genome database. Hg19 46-way alignment scores are not strand-symmetric (https://genome.cshlp.org/content/suppl/2009/10/27/gr.097857.109.DC1/supplement.pdf, S2.6), and minus-strand scores are thus inappropriate to include in transcriptomic analyses and were excluded. For meta-phyloP scores, all scores for the sequence GCAUG were averaged at each base in 3’ UTRs and shallow introns (+250 bases) and normalized to the average phyloP score of the *5*mer.

### Identification of secondary motifs by iterative *k*mer analysis

Initial *5*mer enrichments were determined by generating five-base sequences from the 3’ UTR regions of each valid oligonucleotide in a 0 nM and pulldown experiment. *5*mer enrichment values were then determined by dividing the normalized count of a particular *5*mer in the pulldown by the normalized counts of the *5*mer in the 0 nM experiment. Of the 1024 *5*mers, GCAUG had the highest *R* value. To identify other sequences that influence binding, we took an iterative approach in which all oligonucleotides containing the highest *R* value *5*mer were removed from consideration, and enrichments for all other *5*mers were regenerated from this set. Following identification of the primary motifs (GCAUG, GCACG) in the 1.1 uM experiment, the top *5*mer in each subsequent iteration was considered a secondary motif (GUUUG, GAAUG, UUUUU, UGCAU, AAAAA, GUAUG, GCUUG). After four iterations, there was insufficient power to continue the iterative method with ∼18,000 of ∼64,000 sequences remaining. All remaining *5*mers of the format GNNUG, NGCAU, or W_5_ that had *R* value two standard deviations above the mean were considered secondary motifs (GCCUG, GCCUG, CGCAU, AGCAU, AUAUA, UAUAU).

### eCLIP peak enrichment and iCLIP metaplot analyses

eCLIP enrichment values were produced from significant RBFOX2 eCLIP read peaks in HepG2 cells obtained from the ENCODE Project Consortium^40^. RBFOX2-pulldown peak densities were normalized to a no-protein input control to produce enrichment values roughly analogous to nsRBNS *R* values to facilitate comparisons between the two assays.

RBFOX2 individual nucleotide crosslinking and immunoprecipitation (iCLIP data) from mouse embryonic stem cells^27^ was analyzed at RBFOX secondary motif sites. Adapters and barcodes were trimmed prior to mapping with STAR to the mm10 genome following standard ENCODE guidelines (http://labshare.cshl.edu/shares/gingeraslab/www-data/dobin/STAR/STAR.posix/doc/STARmanual.pdf, page 7). Duplicate PCR reads were removed from the mapped reads to generate final reads. These reads were aligned in a metaplot centering on all possible *5*mers to visualize an iCLIP meta-peak. Peak height was quantified with a CLIP enrichment (CE) score centered on position 1 of the *5*mer. For 3’ UTR peaks, the CE score was calculated as the sum of the read coverage between positions 10 to 15 divided by the sum of the read coverage between positions –85 to –80. For the intronic peaks, the CE score was calculated by the read coverage between positions 5 and 10 divided by the read coverage at positions −85 to −80. The ranges differed between 3’ UTR and intronic peaks due to different maximum peak heights in these regions; the maximum range was chosen for each region. Control scores were produced using an untagged RBFOX2 protein with identical data processing; these background reads were subtracted from the metaplots and CE scores to eliminate iCLIP noise.

### HiTS-CLIP motif enrichment

Two RBFOX1 HiTS-CLIP^8, 14^ datasets, in mouse whole brain and differentiated mature neurons, respectively, were analyzed for the presence of secondary motifs. Both datasets were mapped with STAR after removing duplicate reads following ENCODE guidelines, except for requiring mapped reads to be completely unique (-- outFilterMultimapNmax 1). For the Jacko *et al*.^8^ data, read length was relaxed to accommodate the slightly shorter average HiTS-CLIP read length (--outFilterMatchNminOverLread 0.33). Mapped reads were then processed into CLIP peaks using CLIPper with standard specifications. For each dataset, only peaks shared between two replicates were considered in subsequent analyses. Additionally, only peaks in 3’ UTRs and shallow (splice site-proximal) regions of introns were analyzed. For each CLIPper peak, a region ±50 from the reported peak apex was analyzed for the frequencies of all possible *5*mers to report a total *5*mer frequency for each dataset analyzed. *5*mer frequencies in CLIPper regions were normalized to the total *5*mer frequencies in either 3’ UTRs and shallow introns to generate *5*mer enrichments. To analyze the enrichments of peaks containing primary, primary and secondary, or secondary motifs, a peak with one of these motifs in the 100-base region was considered to contain the motif. To assess signal over background for these groups, artificial 100-base intervals derived from 3’ UTRs and shallow introns, respectively, were analyzed identically and frequencies were compared.

### Splicing reporter assay

We cloned primary and secondary Rbfox motifs GCAUG.1, GCAUG.2, GCUUGx6, GAAUGx6, or GUUUGx6 250 bases downstream of the alternative exon of the RG6 splicing reporter^43^ (see sequences below, with altered nucleotides in capitals) using custom-designed oligonucleotides (IDT) with InFusion cloning (Takara Bio #638920). The GFP of a pEGFP rbFOX1 plasmid (Addgene #63085) was replaced with Cerulean (Cerulean-N1 Addgene #54742) to produce a Cerulean:Rbfox1 vector. The downstream Rbfox1 was also removed to produce a Cerulean:NULL control plasmid. The sequences inserted into each plasmid are listed below:

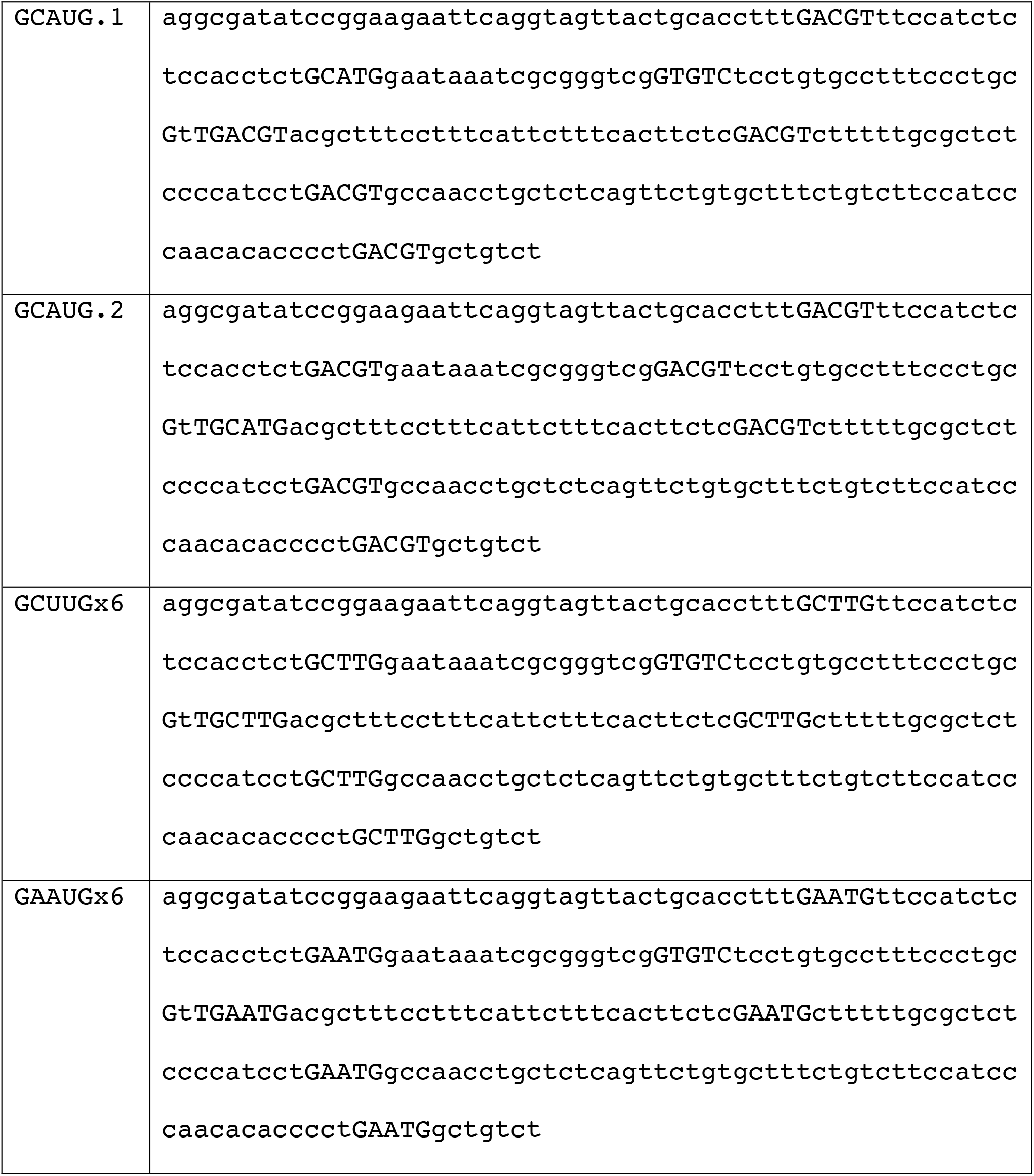

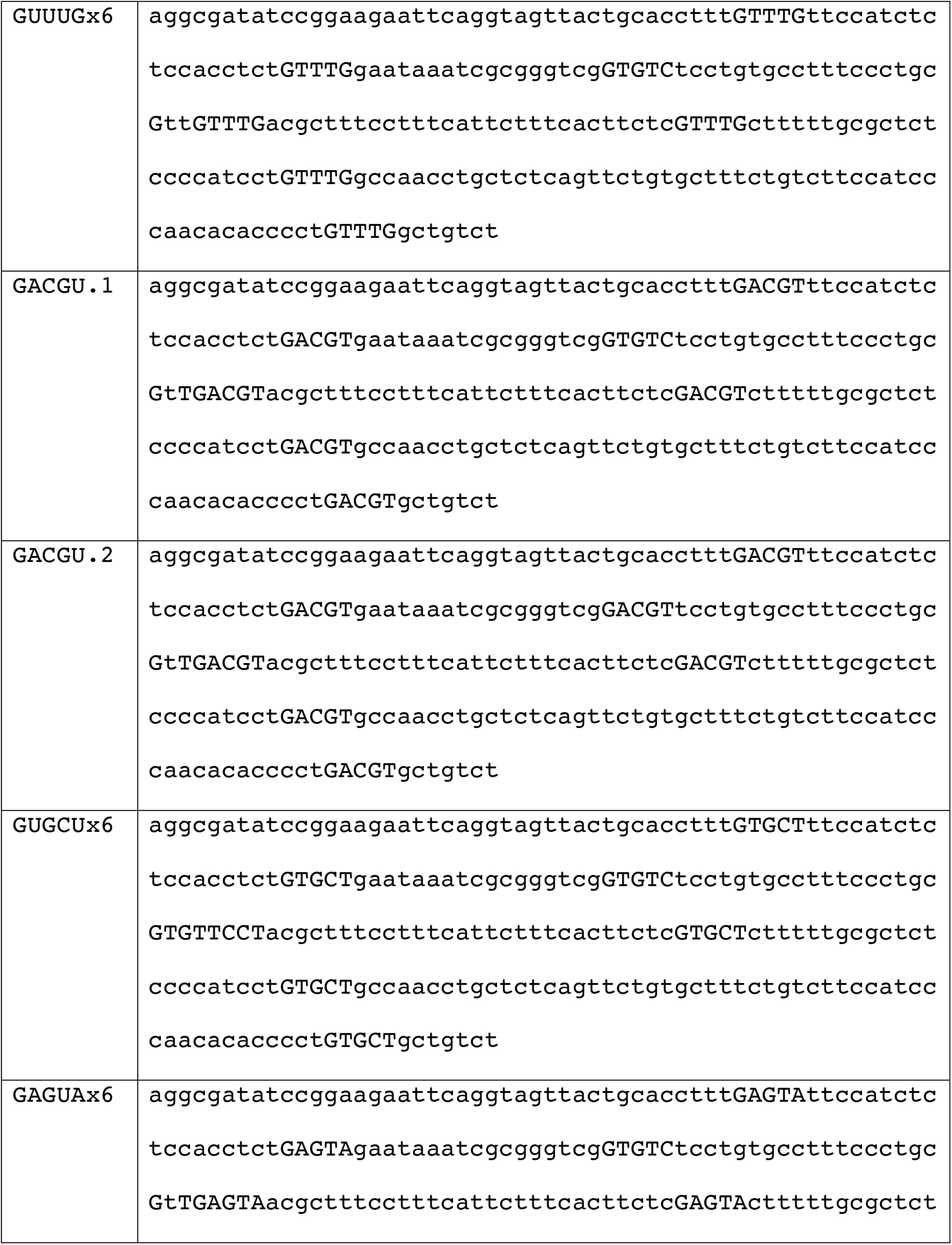

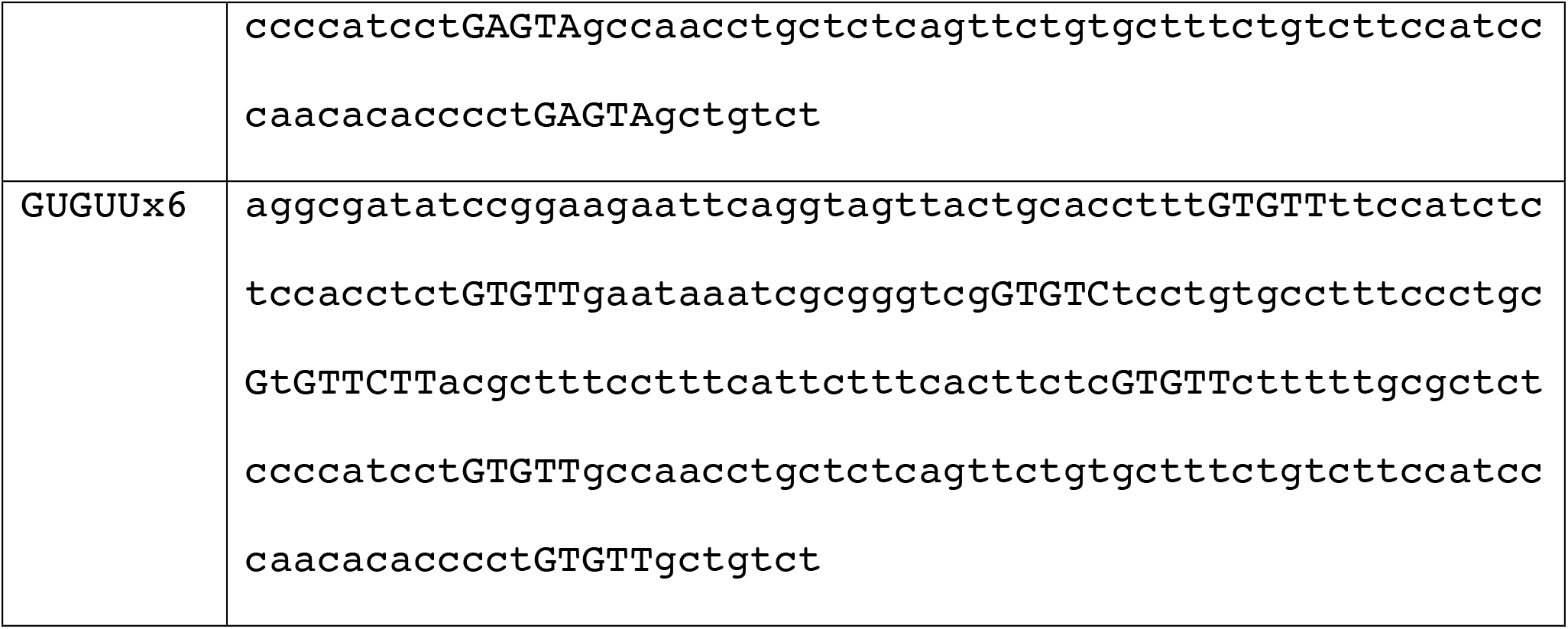

A far-downstream GCAUG endogenous to the plasmid was included in all plasmids. 100 ng of RG6 construct, 300 ng of Cerulean:Rbfox or Cerulean:NULL was transfected into HEK293T cells in a 24-well plate with 1 uL of lipofectamine. Transfected cells were harvested after 24h. The cells were washed twice with 2 mL PBS and RNA was extracted using the Qiagen RNAeasy Kit (#74104). Three replicates of each condition were subjected to fluorescent PCR with a FAM-labelled forward primer (below) and with Phusion polymerase (NEB #M0530S) for 32 cycles. The product was imaged on a Typhoon FLA 9500. Resultant bands were quantified using ImageJ to produce relative Percent Spliced In (PSI) values.

### Flow cytometry and data processing

400 ng of RG6 construct, 400 ng of Cerulean:Rbfox or Cerulean:NULL was transfected into HEK293T cells in a 24-well plate with 4 uL of lipofectamine. Transfected HEK293T cells were harvested after 48 hours of transfection in 6-well plates, with 10^6^ cells per well. After the media was removed, cells were gently washed in 1 mL PBS, and then resuspended into 1 mL ice-cold PBS with 1% BSA and 2 mM EDTA. Cell suspensions were collected into test tubes through a single-cell strainer (Fisher Scientific; Corning Falcon #352235) on ice. Flow cytometry was carried out with the LSR II flow cytometer (BD Biosciences), with the 405 nm laser and 450/50 nm filter for Cerulean, 488 nm laser and 515/20 nm filter for EGFP, and 561 nm laser and 610/20 nm filter for dsRED. A total of 30,000 events from single, live cells were acquired for each treatment set, and processed using the FlowJo software. Minor channel spillover between EGFP and Cerulean was compensated using single-fluorophore controls.

Cerulean signal was normalized by dividing over the median Cerulean signal of the no-plasmid control, to account for background fluorescence. To only include cells with at least one copy of both plasmids transfected, we used the 99^th^ percentile of the signal from the three respective fluorophores in the no-plasmid control as the thresholds, and filtered for events with Cerulean above threshold, and with EGFP and/or dsRED above threshold. Events with EGFP or Cerulean signal above 10^4.5^, or dsRED signal above 10^3.2^, were discarded due to signal anomaly near the detector saturation limit. Events were sorted into bins by log_2_-transformed normalized Cerulean signal, dividing at 2.2, 3.0, 4.5, 6.0, and 7.5, to obtain six bins with generally similar number of events each. Log_2_-transformed ratio of EGFP signal to dsRED signal was used as the readout for the splicing ratio.

### Neuronal differentiation analysis

Using a deep transcriptomic sequencing dataset characterizing neuronal differentiation from mouse embryonic stem cells (mESCs) to glutamatergic neurons^39^, we examined the use of Rbfox primary and secondary motifs in neuronal differentiation. Each of eight (ESC, NESC, RG, DS1, DS3, MAT16, MAT21, MAT28) time points was analyzed for total Rbfox (Rbfox1, Rbfox2, Rbfox3) expression using kallisto^56^ with standard parameters. Splicing events were analyzed with rMATS.4.0.2^57^ using standard specifications between the radial glia (RG) stage and all subsequent events (e.g. RG–DS1, RG–DS3, RG–MAT16, etc.) as well as between each interval (e.g. ESC–NESC, NESC–RG, RG–DS1, etc.). For all significantly changing cassette exons (FDR < 0.1), the secondary motif content of the first 250 bases of the downstream intron was computed and correlated with the magnitude of the inclusion of the upstream exon as reported by rMATS.

### Gene Ontology

Genes regulated *exclusively* by primary motifs (primary motif-mediated) or secondary motifs (secondary motif-mediated) in the 250 nt of the intron downstream of an exon increasing in inclusion from DS1 to DS3 were subjected to Gene Ontology analysis with GOrilla^58, 59^. Results were then filtered by FDR < 0.1, B > 99, and b > 9. Background genes were all genes expressed >1 transcript per million (tpm) in DS3 as assessed by kallisto^56^.

### Stability analysis

We analyzed data from Lee et al.^12^ in which *Rbfox1* and *Rbfox3* siRNA knockdown in mouse hippocampal neurons were rescued with the cytoplasmic isoform of Rbfox1. The 3’ UTR *5*mer conservation of transcripts that were destabilized upon Rbfox knockdown and rescued by the cytoplasmic isoform of Rbfox1 were assessed using phyloP scores. Secondary motif *5*mers of interest were compared to all other *5*mers contained in the same 3’ UTRs.

### Linear regression analysis of motif frequency and alternative splicing

Starting from a table with PSI values for 1,909 alternative exons regulated in neuronal cells and a list of RBP expression values (courtesy of the Chaolin Zhang lab, Columbia University), underlying data^4^, cell types were grouped by the sum of Rbfox1, Rbfox2, and Rbfox3 expression values. For each group of cell types, the arithmetic mean of PSI values was computed, from which followed a ΔPSI value for changes in exon inclusion between Low and Medium, Medium and High, and Medium and Highest Rbfox-expressing cells. Next, for each alternative exon, the intronic sequences 8nt to 250 nt downstream of the exon were scanned for the presence of Rbfox motif *5*mers. Then, for each vector of ΔPSI values across all exons, linear regression was performed with the number of Rbfox motif occurrences as explanatory variable (using the python scipy.stats.linregress function of the scipy package^60^). The resulting P-values and *r* values were plotted using matplotlib^61^.

### RBNS calibration to surface plasmon resonance (SPR) database

We computed RBFOX2 and RBFOX3 7-mer enrichments from RNA Bind-n-Seq (RBNS) experiments performed with 1.1 and 1.3 μΜ respectively^26, 28^. These *R*-values should directly track with the occupancy of RBFOX protein, but also contain a contribution from non-specific sequences, either captured through the apparatus or due to the fact that 20 nt and 40 nt random sequences were used (and not *7*mers, see Lambert *et al*. 2014^28^ for a discussion). We therefore estimated the *R*-value of non-specific *7*mers R_ns_ from the bottom percentile of R-values and derived corrected R-values as R’ = R + R_ns_ * (R – 1)/(1 – R_ns_). The corrected R-values display a higher dynamic range and provide a slightly better fit to SPR reference affinities (not shown). The results change only marginally if R_ns_ is estimated from other percentiles (5 or 10, not shown).

Next, we compiled a list of dissociation constants for 22 *7*mer sequences binding to RBFOX1, measured via SPR from Auweter *et al*.^41^ and Stoltz^29^. We then used the scipy.stats.linregress function to find an optimal linear relationship between log(R’) and log(K_d_) for these sequences. We then extended the linear interpolation to all *7*mer R’ values to assign approximate dissociation constants. The two experimental replicates (with RBFOX2 and RBFOX3, using 20 nt and 40 nt random sequences) yielded highly correlated results (Supplementary Figure S10c). We then computed average *5*mer dissociation constants as K_d_ ^5^ = exp(<log(K_d_ ^7^)>), where < … > denotes the arithmetic mean over all *7*mers containing the *5*mer of interest, and across both calibrated data sets for RBFOX2 and RBFOX3. Values obtained by subtracting or adding one standard error of the mean were used to estimate the error bars. In the affinity histogram (Supplementary Figure 10d), non-primary or secondary *5*mers containing partial primary motifs (GCA, AUG, and ACG) were masked, because RBNS *R* values are always contaminated by the enrichment of *k*mers that overlap authentic high affinity motifs but are not necessarily directly bound (see Lambert et al. 2014^28^).

### Estimation of the intronic sequence content of the nucleus

Common estimates for total mRNA copy numbers per cell range from 100,000 to 1,000,000 molecules per cell (consistent with 0.1 to 1 pg of mRNA per cell). To arrive at a rough estimate of how much intronic RNA is made, we therefore convert the estimated average half-life time of mRNA molecules of T_h_ = 5 to 10 hours into a rate k_deg_ = log(2)/T_h_ and assume that mRNA decay is balanced by new mRNA production. We considered two scenarios that span the anticipated range of RNA concentrations: a small cell (concentrated RNA) scenario with 100,000 mRNAs in a cell volume of 500 μm^3^ with T_h_ = 5 h; and a large cell (dilute RNA) scenario with 1,000,000 mRNAs in a cell volume of 10,000 μm^3^ with T_h_ = 10 h. These estimates were based on values in BioNumbers.org (cell volume ID 112112, ID 106320, mRNA copy number ID 111220, half-life times ID 106378, ID 104747). We further assume that the nucleus comprises ∼10% of the cell’s volume and that effectively 70% of that volume are available for the diffusion of intronic RNA and Rbfox proteins (subtracting nucleolus and chromatin). Our findings are not dependent on these exact values.

By iterating over the catalog of mouse transcripts (gencode M97) and weighting each encountered intron with the RNA-seq-derived tpm (transcripts per million) expression value for the harboring transcript (using kallisto^56^) we thus estimate the amount of intronic sequence being transcribed per minute. Assuming an average intronic half-life time of 1 minute, we conclude that, on average a mouse neuronal cell nucleus may harbor between 1.2 and 6 million intronic *5*mer sequences that could represent potential protein binding sites.

Using the same tpm-weighting scheme, we then compute the share of this total nuclear, intronic *5*mer concentration allotted to each *5*mer.

### Estimation of nuclear Rbfox protein concentration

With a given estimate of total cellular mRNA copy number, tpm (transcripts per million) values correspond directly to cellular mRNA copies/cell. To estimate how many protein molecules are present per mRNA copy, we investigated Schwanhäusser et al. 2011^45^ who report ∼10,800 proteins/mRNA for *Rbm9* (aka *Rbfox1*) in NIH 3T3 cells and a median of ∼2,800 proteins/mRNA. Li et al. 2014^62^ (BioNumbers.org ID 110236) suggest that this might be an underestimate and place the median at 9,800 proteins/mRNA. We therefore consider the range between 2,800 and 10,000 as reasonable Rbfox protein to mRNA ratio, while assuming that the bulk of Rbfox protein is nuclear. This range reflects the shaded areas in Figure 6c corresponding to low Rbfox expression (10 TPM) and the highest Rbfox expression (1,900 TPM) observed among the neuronal cell types.

### Equilibrium model for protein binding to a diverse pool of binding sites

We follow the procedure employed in Jens and Rajewsky^44^ for miRNA binding sites. Briefly, in equilibrium, assuming a well-mixed compartment, every potential binding site interacts with the same pool of free (unbound) protein [P_free_]. Further, each binding site’s occupancy is assumed to be solely dependent on its primary sequence affinity, represented by a *5*mer (no cooperativity, no competition with other proteins or RNA structure). Under these assumptions, we can compute the occupancy for each *5*mer i using its SPR-calibrated, RBNS-derived K_d_ (see above) as O_i_ = [P_free_] / ([P_free_] + K_d_). At the same time, mass-action dictates that [P_free_] = [P_total_] – [P_bound_]. And [P_bound_] = Σ_i_ O_i_ * c_i_ (where c_i_ is the estimated concentration of *5*mer i in the nucleus). We numerically find [P_free_] to satisfy these constraints to find the occupancy of Rbfox motifs and the fraction of Rbfox protein alotted to these sites as f = Σ_j_ O_j_ * c_j_ / [P_total_] (where j are motif *k*mers). Non-primary, non-secondary *5*mers with partial overlap to primary motifs (139 out of 1,024 *5*mers) were ignored for this analysis.

## Acknowledgments

We thank past and present members of the Burge Lab, John Conboy, and Inga Jarmoskaite for helpful comments on the manuscript. We gratefully acknowledge the courtesy of the Chaolin Zhang lab of Columbia University, who shared intermediate results from Weyn-Vanhentenryck *et al*.^4^ used for Figure 6 (gene expression, PSI values, exon coordinates). M.J. received an EMBO Long Term Fellowship ALTF-1130-2015.

## Author contributions

B.E.B. performed experiments, analyzed data, and wrote the manuscript. M.J. analyzed data and wrote the manuscript. P.Y.W. performed experiments. C.B.B. supervised the study and wrote the manuscript.

## Competing interests statement

The authors declare no competing interests.

