## Supplemental Figures for "Secondary motifs enable concentration-dependent regulation by Rbfox family proteins"

Supplemental Information.

#### Figure S1

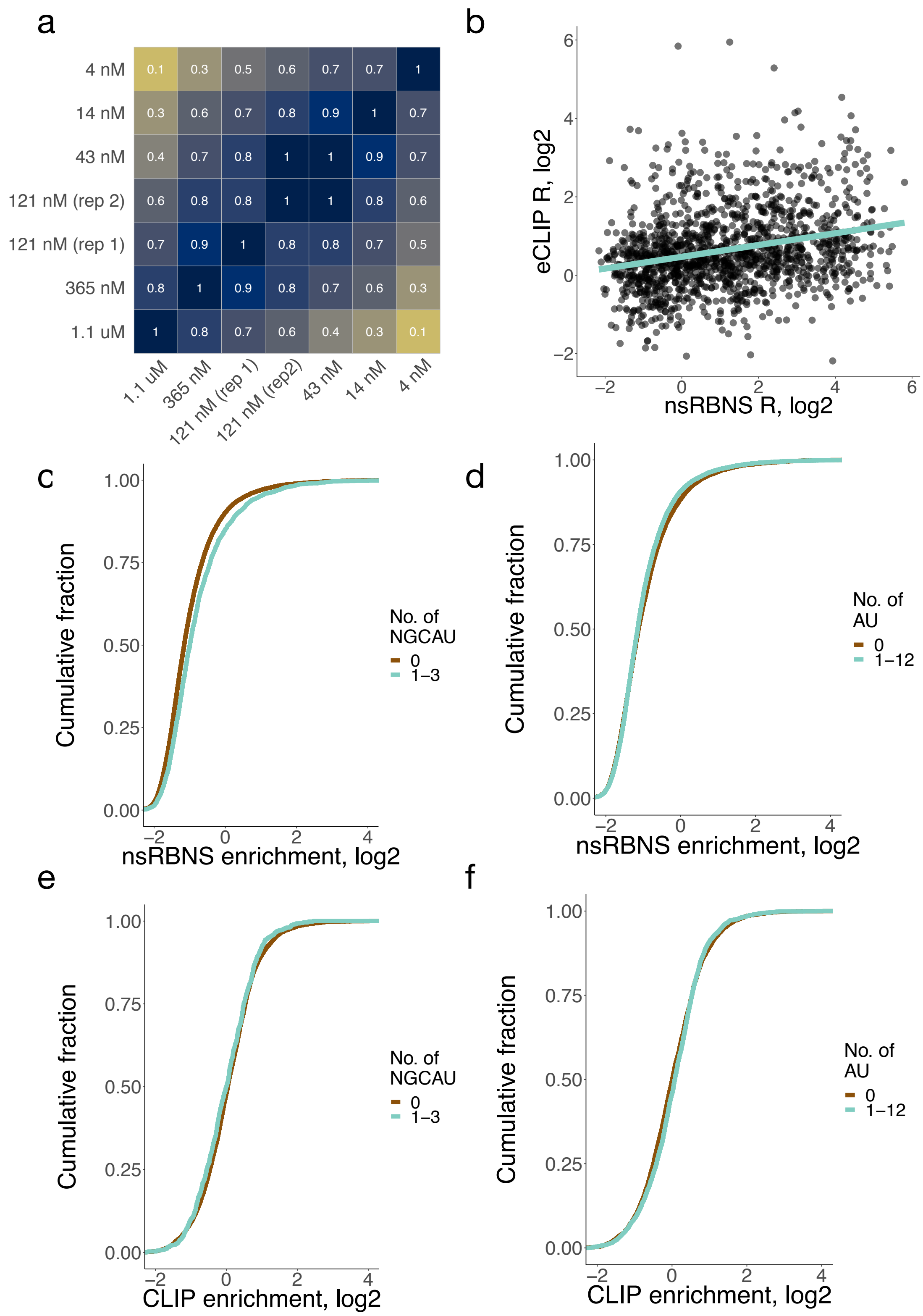
a
4 nM
14 nM
43 nM
121 nM (rep 2)
121 nM (rep 1)
365 nM
1.1 uM
1.1 uM
365 nM
121 nM (rep 1)
121 nM (rep2)
43 nM
14 nM
4 nM
b
eCLIP R, log2
nsRBNS R, log2
c
Cumulative fraction
nsRBNS enrichment, log2
No. of NGCAU
0
1–3
d
Cumulative fraction
nsRBNS enrichment, log2
No. of AU
0
1–12
e
Cumulative fraction
CLIP enrichment, log2
No. of NGCAU
0
1–3
f
Cumulative fraction
CLIP enrichment, log2
No. of AU
0
1–12

**Figure S1. RBFOX2 nsRBNS reveals binding to a set of moderate-affinity secondary motifs.** a. Correlations among seven natural sequence nsRBNS experiments. Pearson correlations are reported for any sequence with an enrichment (*R*) value greater than 1. Darker color indicates a higher correlation. b. Correlation of nsRBNS *R* with eCLIP enrichment at oligo-derived regions for all oligonucleotides or sequence regions containing a single GCAUG Rbfox primary motif. c. *R* value distribution of nsRBNS sequences containing increasing numbers of NGCAU motifs. d. *R* value distribution of nsRBNS sequences containing increasing numbers of AU motifs. e. RBFOX2 eCLIP in HepG2 at library positions in the transcriptome for NGCAU motifs. RBFOX2 peaks were compared to an IgG control to determine enrichments. f. RBFOX2 eCLIP in HepG2 at library positions in the transcriptome for AU motifs. RBFOX2 peaks were compared to an IgG control to determine enrichments.

### Figure S2

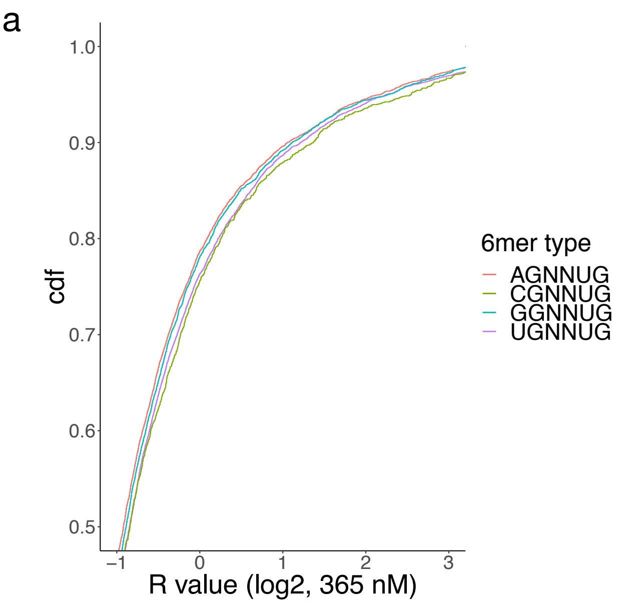
a
1.0
0.9
0.8
0.7
0.6
0.5
cdf
–1
0
1
2
3
R value (log2, 365 nM)
6mer type
AGNNUG
CGNNUG
GGNNUG
UGNNUG

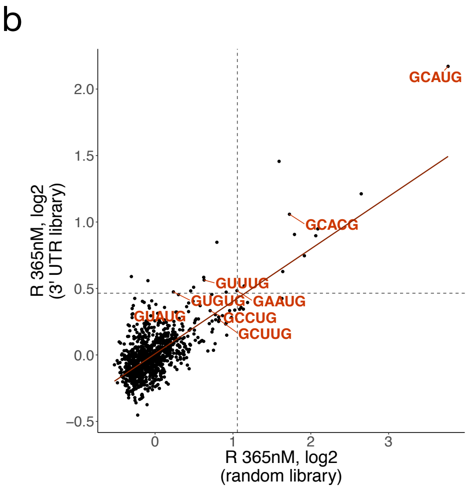
b
GCAUG
GCACG
GUUUG
GUGUG
GAAUG
GUAUG
GCCUG
GCUUG
2.0
1.5
1.0
0.5
0.0
–0.5
R 365nM, log2
(3' UTR library)
0
1
2
3
R 365nM, log2
(random library)

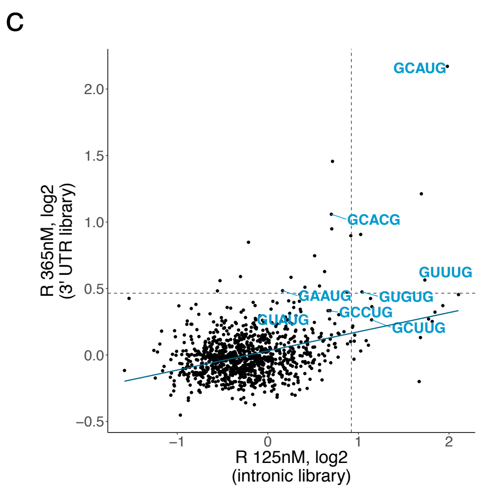
c
GCAUG
GCACG
GUUUG
GAAUG
GUGUG
GCCUG
GUAUG
GCUUG
2.0
1.5
1.0
0.5
0.0
–0.5
R 365nM, log2
(3' UTR library)
–1
0
1
2
R 125nM, log2
(intronic library)

**Figure S2. Different nsRBNS libraries emphasize different *5*mer binding preferences for RBFOX2.** a. *R* value distribution of nsRBNS sequences containing exactly two copies of different *6*mer classes. b-c. Comparison of random (b) and intronic natural sequence (c) RBNS with 3' UTR nsRBNS *5*mer enrichments. Primary and secondary motifs are labelled in red and blue, respectively. Dotted lines show 2.5 standard deviations above the mean.

### Figure S3

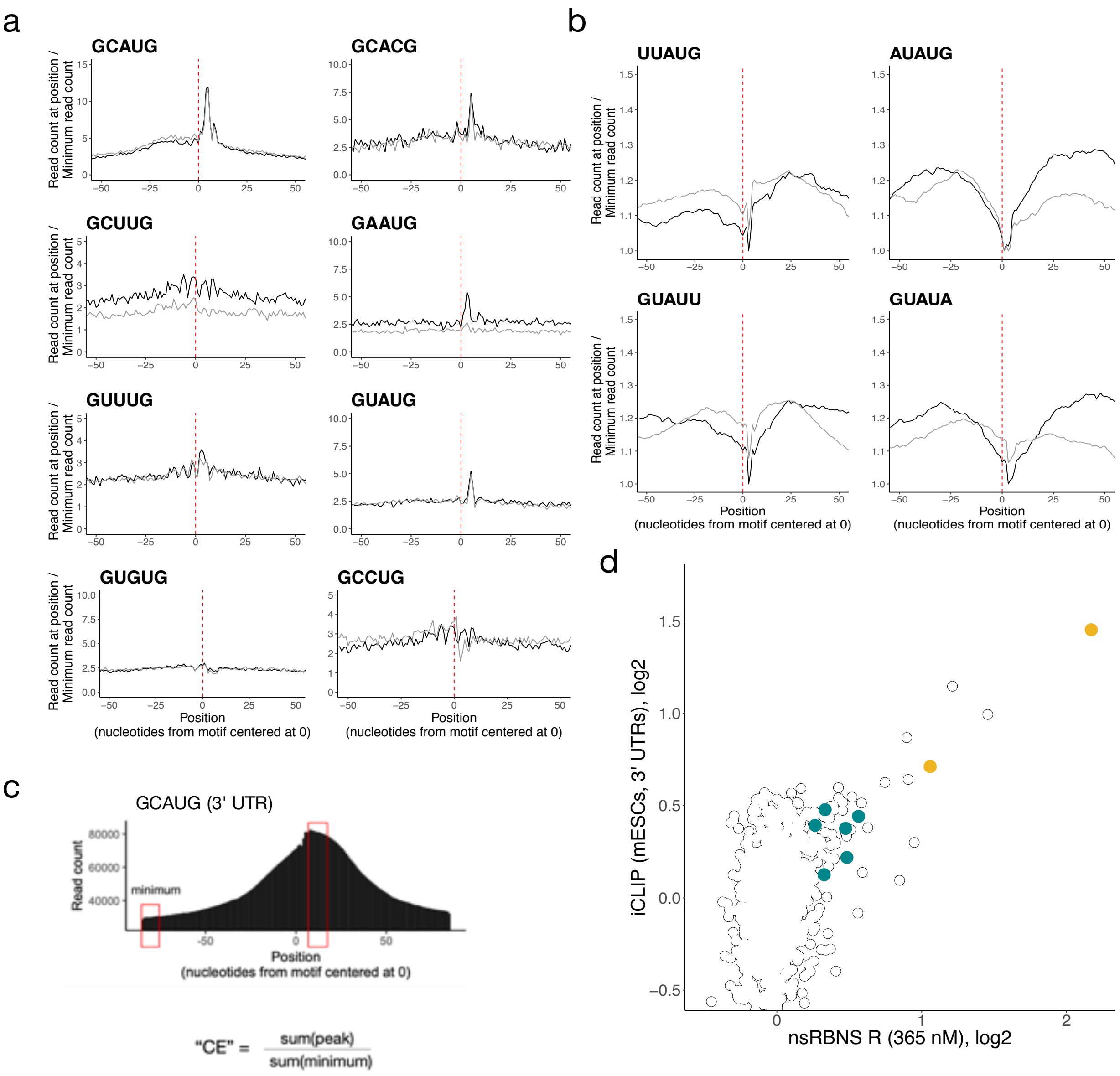

a
GCAUG
GCACG
GCUUG
GAAUG
GUUUG
GUAUG
GUGUG
GCCUG
Read count at position / Minimum read count
Position (nucleotides from motif centered at 0)
b
UUAUG
AUAUG
GUAUU
GUAUA
Read count at position / Minimum read count
Position (nucleotides from motif centered at 0)
c
GCAUG (3' UTR)
Read count
minimum
Position (nucleotides from motif centered at 0)
"CE" = sum(peak) / sum(minimum)
d
iCLIP (mESCs, 3' UTRs), log2
nsRBNS R (365 nM), log2

**Figure S3. RBFOX2 iCLIP demonstrates broad agreement with nsRBNS.** a. Some secondary motifs show sharp peaks near 0 in a metaplot centered at the motif in introns (black) and 3' UTRs (grey) in RBFOX2 iCLIP data[27]. 5' ends of iCLIP reads containing the motif of interest were aligned with position one of the pentamer at 0 and normalized to the minimum read count in an 80-nt window (50-nt window shown). Y-axis range was reduced for secondary motifs. b. AU-rich nsRBNS motifs do not show characteristic read peaks near 0 in a metaplot centered at the motif in introns (black) and 3' UTRs (grey) in RBFOX2 iCLIP data[27]. iCLIP reads containing the motif of interest were aligned with position one of the pentamer at 0 and normalized to the minimum read count in an 80-nt window (50-nt window shown). Y-axis range was reduced for secondary motifs. c. Schematic showing the generation of a clip enrichment (CE) score from iCLIP data. After generation of a metaplot, the read count at the peak apex was divided by the read count at its lowest point to generate a CE score analogous to an enrichment. d. Correlation of iCLIP- and nsRBNS-enriched 5mers in 3' UTRs. CLIP enrichment (CE) scores were computed for iCLIP peaks. Secondary motifs indicated in teal, primary motifs indicated in gold.

### Figure S4

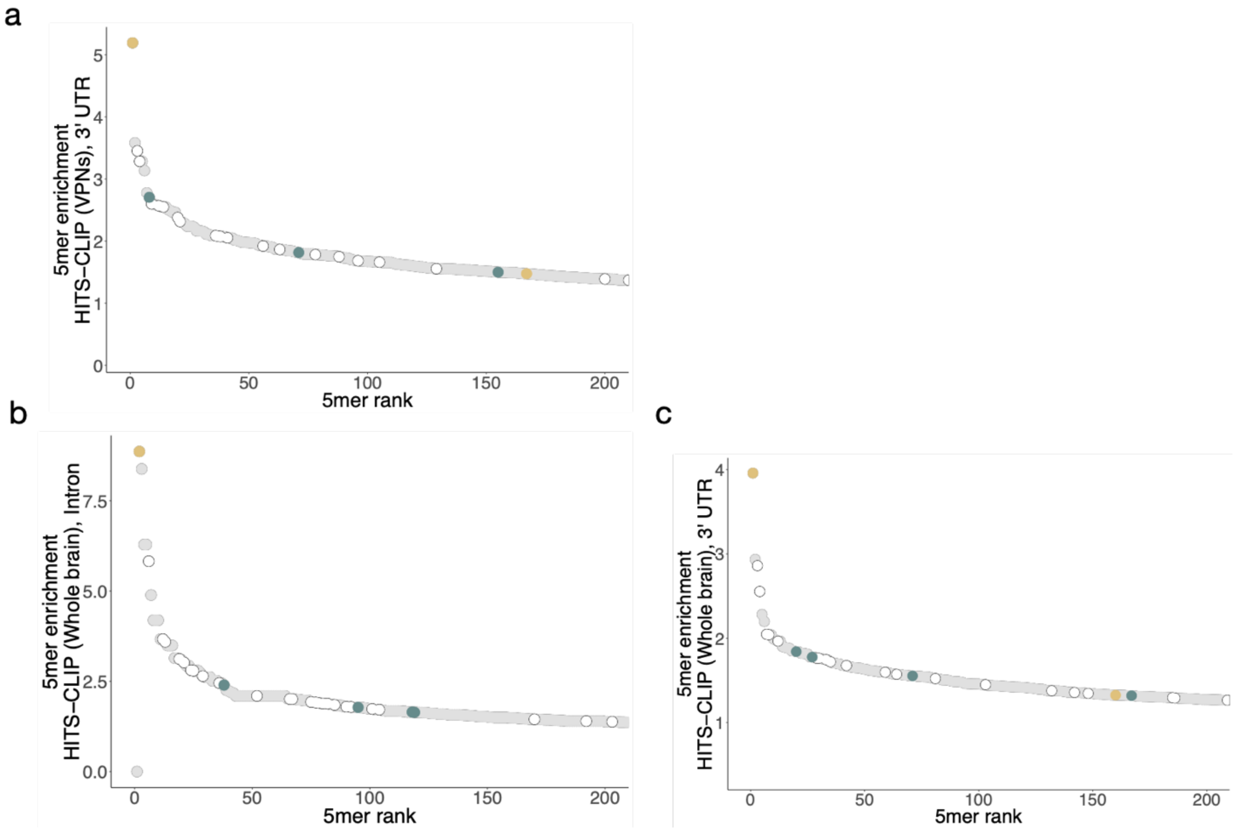
a
5mer enrichment
HITS-CLIP (VPNs), 3' UTR
0
1
2
3
4
5
0
50
100
150
200
5mer rank
b
5mer enrichment
HITS-CLIP (Whole brain), Intron
0.0
2.5
5.0
7.5
0
50
100
150
200
5mer rank
c
5mer enrichment
HITS-CLIP (Whole brain), 3' UTR
1
2
3
4
0
50
100
150
200
5mer rank

**Figure S4. Enrichment of 5mers in HiTS-CLIP.** 5mer enrichment of top 200 5mers in two HiTS-CLIP datasets in both introns and 3' UTRs. 5mer enrichment was calculated by determining the frequencies of all 1,024 5mers in CLIP peaks in each region and dataset and subsequently normalizing to control peaks from that region. Peaks from (a) Mouse ventral spinal neuron 3' UTR HiTS-CLIP, (b) Mouse whole brain intronic HiTS-CLIP, and (c) Mouse whole brain 3' UTR HiTS-CLIP were analyzed. Gold indicates primary motifs, teal indicates secondary motifs.

#### Figure S5

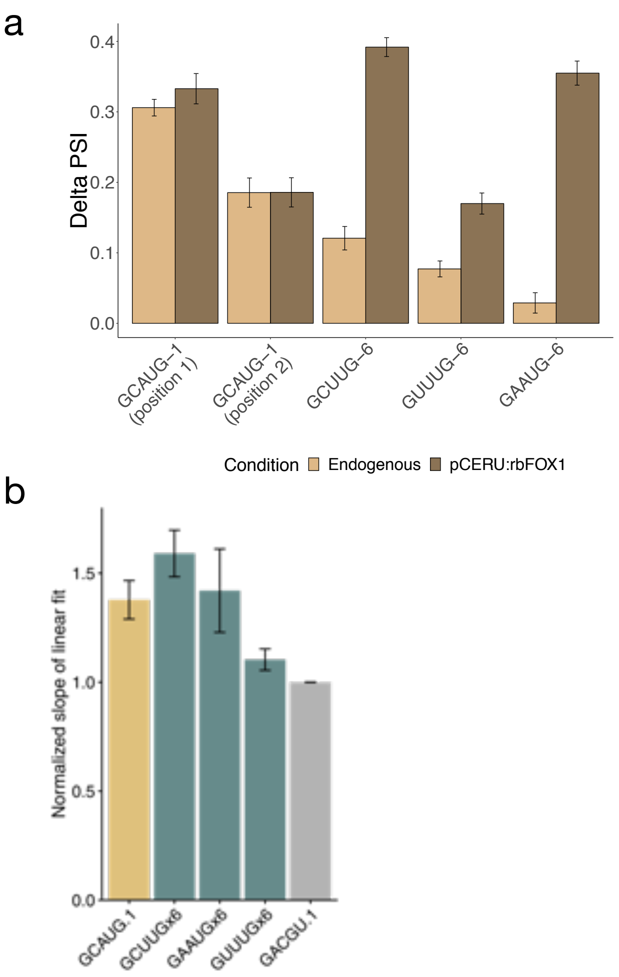

a
0.4
0.3
0.2
0.1
0.0
Delta PSI
GCAUG–1 (position 1)
GCAUG–1 (position 2)
GCUUG–6
GUUUG–6
GAAUG–6
Condition
Endogenous
pCERU:rbFOX1
b
1.5
1.0
0.5
0.0
Normalized slope of linear fit
GCAUG.1
GCUUGx6
GAAUGx6
GUUUGx6
GACGU.1

**Figure S5. Secondary motifs promote inclusion in a splicing reporter in an RBFOX1-dependent manner.** a. Delta PSI values for primary or secondary motif-regulated plasmids in the presence and absence of exogenous RBFOX1. Delta PSI values were computed by subtracting the PSI of the motif minus the PSI of its permuted motif and compared in endogenous versus Rbfox overexpression conditions. b. The slope of linear fit of two flow cytometry replicates were null-subtracted and normalized to their permuted controls. Error bars represent standard error of the mean (SEM).

#### Figure S6a

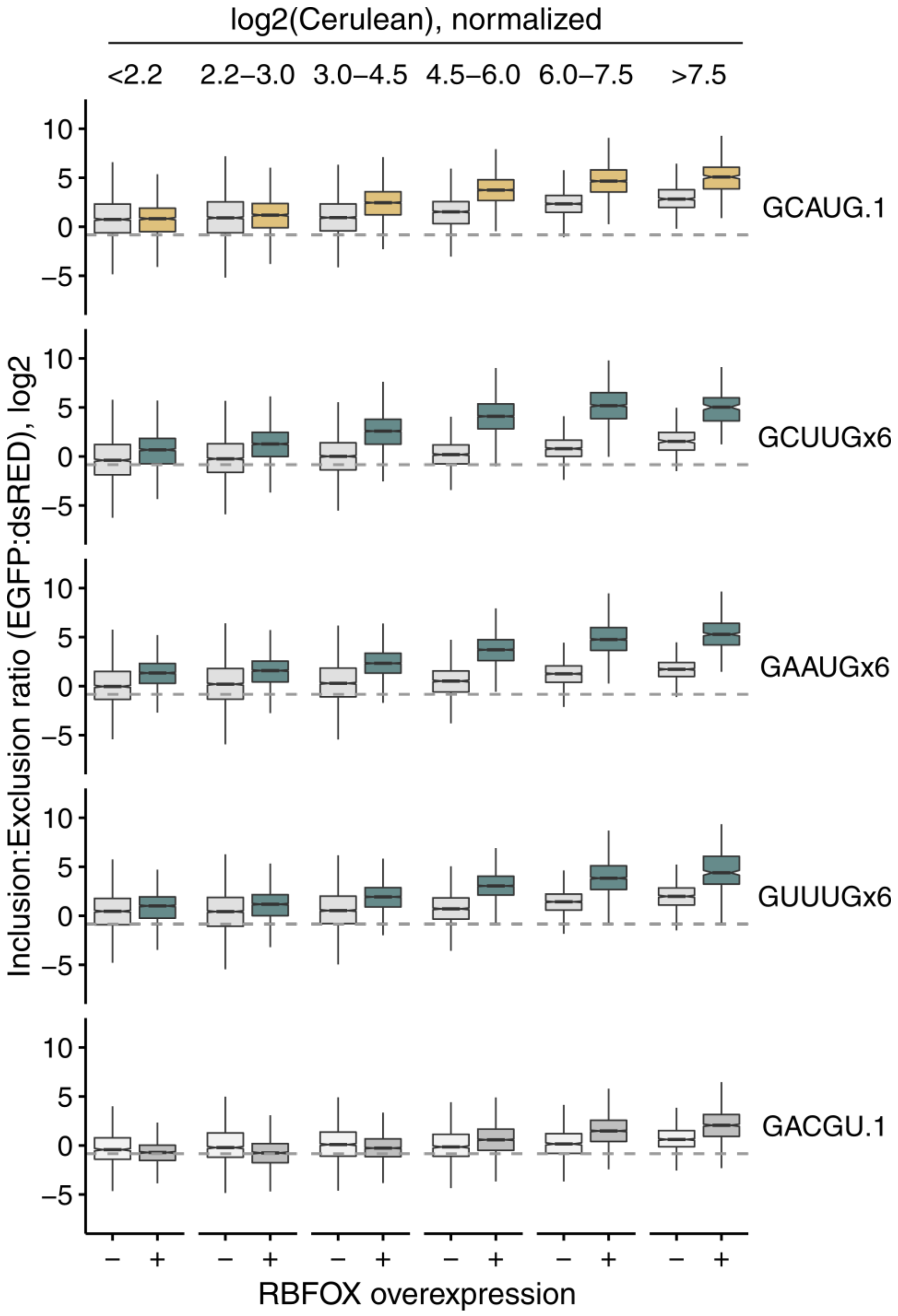
log2(Cerulean), normalized
<2.2
2.2–3.0
3.0–4.5
4.5–6.0
6.0–7.5
>7.5
10
5
0
–5
GCAUG.1
GCUUGx6
GAAUGx6
GUUUGx6
GACGU.1
Inclusion:Exclusion ratio (EGFP:dsRED), log2
–
+
RBFOX overexpression

**Figure S6b**

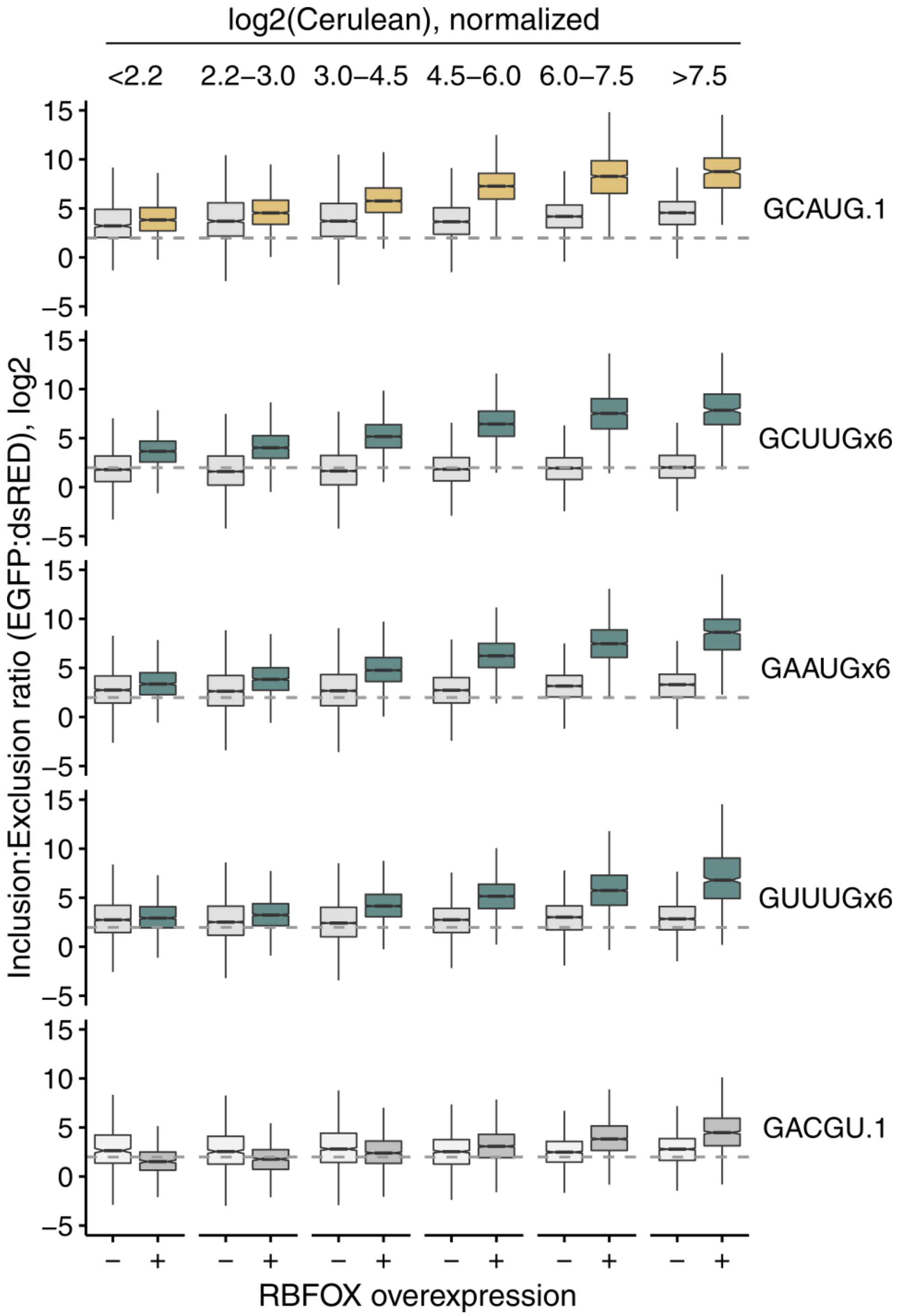

log2(Cerulean), normalized
<2.2
2.2–3.0
3.0–4.5
4.5–6.0
6.0–7.5
>7.5
Inclusion:Exclusion ratio (EGFP:dsRED), log2
15
10
5
0
–5
GCAUG.1
GCUUGx6
GAAUGx6
GUUUGx6
GACGU.1
– +
RBFOX overexpression

**Figure S6c**

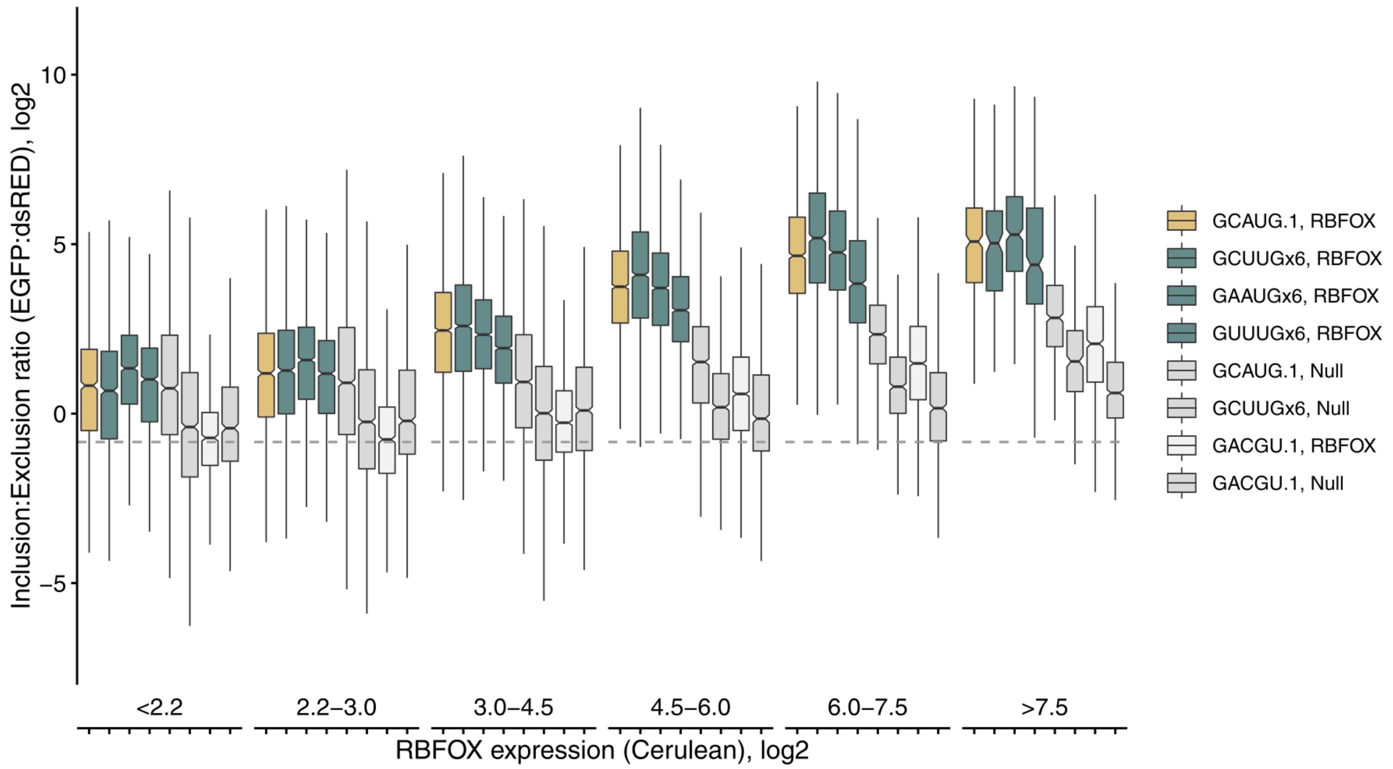

Inclusion:Exclusion ratio (EGFP:dsRED), log2
10
5
0
−5
<2.2
2.2–3.0
3.0–4.5
4.5–6.0
6.0–7.5
>7.5
RBFOX expression (Cerulean), log2
GCAUG.1, RBFOX
GCUUGx6, RBFOX
GAAUGx6, RBFOX
GUUUGx6, RBFOX
GCAUG.1, Null
GCUUGx6, Null
GACGU.1, RBFOX
GACGU.1, Null

**Figure S6. Secondary motifs promote inclusion in a splicing reporter in an RBFOX1-dependent manner at the protein level.** a-b. RG6 plasmids containing one primary motif or six secondary motifs in the presence (gold, teal), or absence (light grey) of Rbfox were co-transfected in HEK293T cells with fluorescently labelled Rbfox1 and monitored by flow cytometry for the inclusion isoform (GFP), exclusion isoform (dsRED), and Rbfox1 (Cerulean) expression at the single-cell level. Controls including a scrambled motif co-transfected with Rbfox1 (dark grey). Two replicates (a and b). c. Six secondary motifs approximate the exon inclusion of one primary motif in an Rbfox1-dependent manner at the protein level, replicate 2. RG6 plasmids containing one primary motif or six secondary motifs were co-transfected in HEK293T cells with fluorescently labelled Rbfox1 and monitored by flow cytometry for the inclusion isoform (GFP), exclusion isoform (dsRED), and Rbfox1 (Cerulean) expression at the single-cell level. Controls including a scrambled motif co-transfected with Rbfox1 (light grey) and scrambled and intact motifs without Rbfox1 (grey) are also shown.

### Figure S7

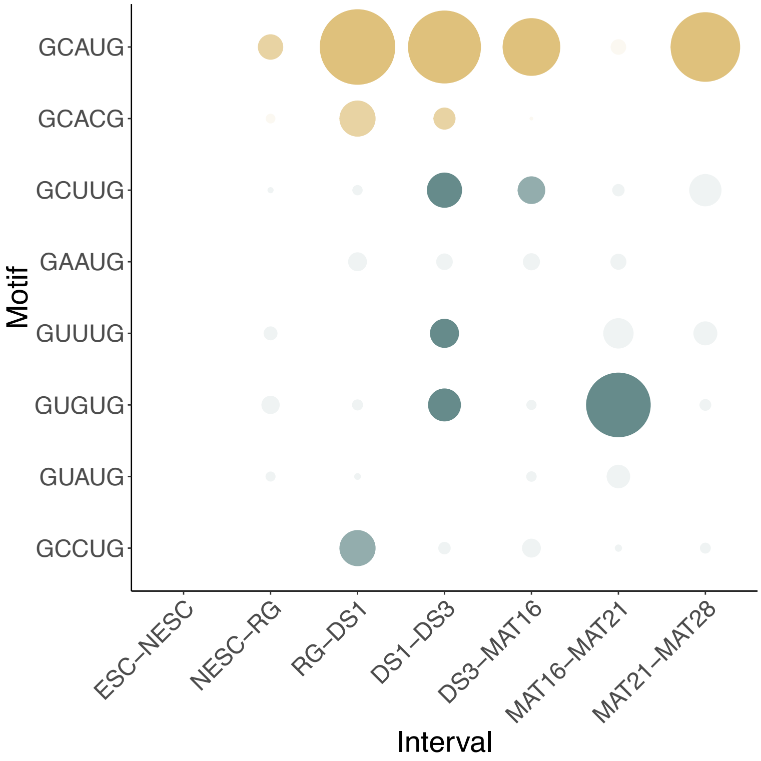
GCAUG
GCACG
GCUUG
GAAUG
GUUUG
GUGUG
GUAUG
GCCUG
Motif
ESC–NESC
NESC–RG
RG–DS1
DS1–DS3
DS3–MAT16
MAT16–MAT21
MAT21–MAT28
Interval

**Figure S7. Secondary motifs become engaged at specific intervals of neuronal differentiation.** Pearson correlation of secondary motif presence with exon inclusion at intervals of neuronal differentiation beginning with embryonic stem cells and progressing to mature 28-day glutamatergic neurons. Size of point indicates correlation coefficient, intensity indicates p-value $< 0.05$.

### Figure S8

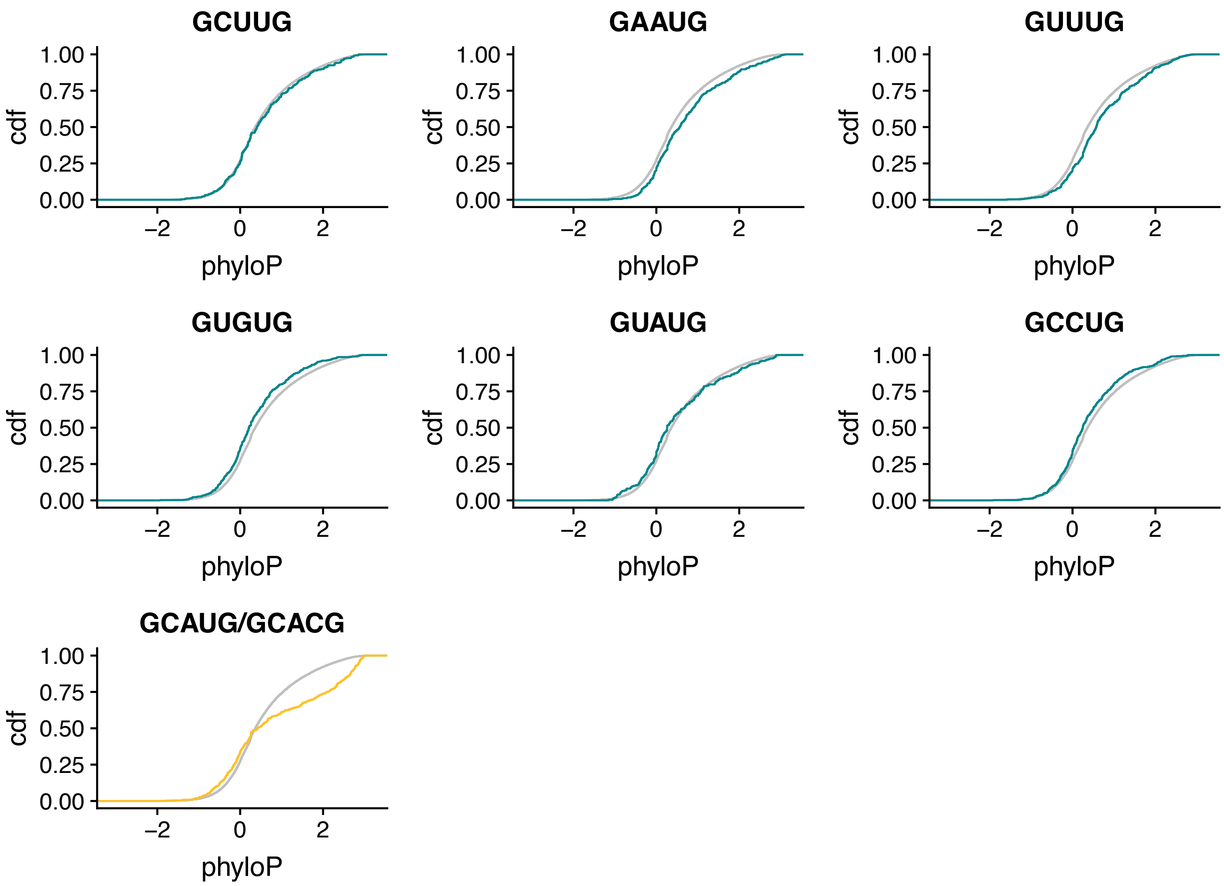

GCUUG
GAAUG
GUUUG
GUGUG
GUAUG
GCCUG
GCAUG/GCACG
cdf
1.00
0.75
0.50
0.25
0.00
−2
0
2
phyloP

**Figure S8. Secondary motifs are more conserved in Rbfox-regulated transcripts.** 3' UTRs of transcripts destabilized by Rbfox1/3 siRNA knockdown and stabilized by its reintroduction were analyzed for 5mer conservation of primary (gold) and secondary (teal) motifs. GAAUG, GUUUG, and GCAYG are significantly ($P < 10^{-3}$) more conserved by Wilcoxon Rank-Sum test, while GUGUG and GCCUG are significantly less conserved by Wilcoxon Rank-Sum. GCUUG and GUAUG were n.s.

### Figure S9

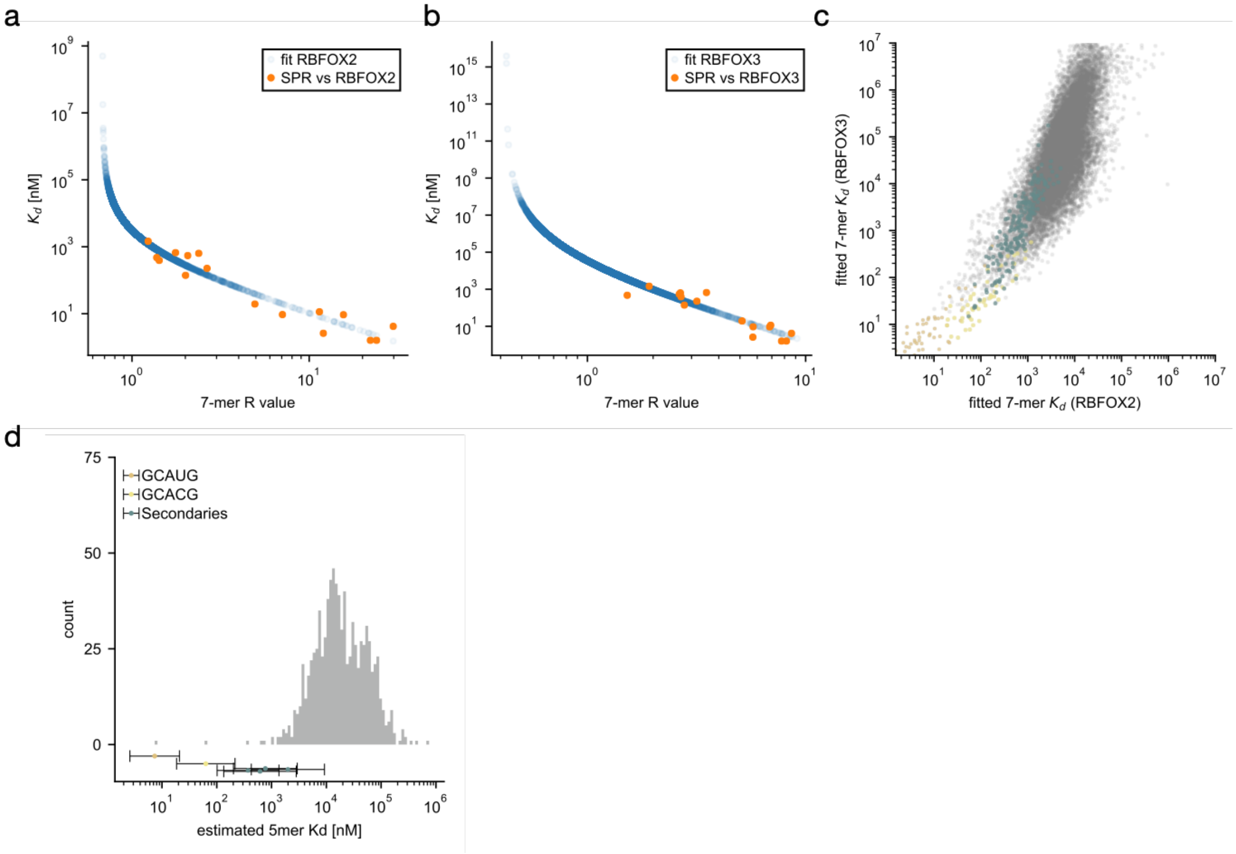

a
fit RBFOX2
SPR vs RBFOX2
$K_d$ [nM]
7-mer R value
b
fit RBFOX3
SPR vs RBFOX3
$K_d$ [nM]
7-mer R value
c
fitted 7-mer $K_d$ (RBFOX3)
fitted 7-mer $K_d$ (RBFOX2)
d
GCAUG
GCACG
Secondaries
count
estimated 5mer Kd [nM]

**Figure S9. Affinity estimation of RBfox secondary motifs.** RBNS 7-mer enrichments (R-value) for 1.1 μM RBFOX2 (a) and 1.3 μM RBFOX3 (b) binding were first corrected for non-specific contributions (R' see Methods) and then linearly correlated with known dissociation constants ($K_d$) for RBFOX1 binding[1,2]. Correlation coefficients between log(R') and log($K_d$) were r=-0.955, P-value=8.379 x $10^{-9}$ (a) and r=-0.915, P-value=6.7 x $10^{-7}$ (b). Scatter plots show estimated $K_d$ as a function of the original, uncorrected R-value. Resulting 7-mer $K_d$ estimates were highly correlated between RBFOX2 and RBFOX3 (c) with r=0.763, P-value ≈ 0. Data for all 7-mers are shown on a logarithmic scale. Primary motif containing 7-mers are highlighted in gold (GCAUG), yellow (GCACG), and teal (secondary motifs GCUUG, GAAUG, GUUUG, GUGUG, GUAUG, GCCUG). Grouping 7-mers by their 5-mer content allows to estimate average $K_d$s for each 5-mer (see Methods). A histogram of these 5-mer dissociation constants is shown in (d), with primary and secondary motifs highlighted as in (c). Motifs GCUUG, GAAUG and GUUUG were considered strong motifs. 136 non-primary or secondary 5-mers with partial overlap to primary motifs GCAUG, GCACG were excluded.

### Figure S10

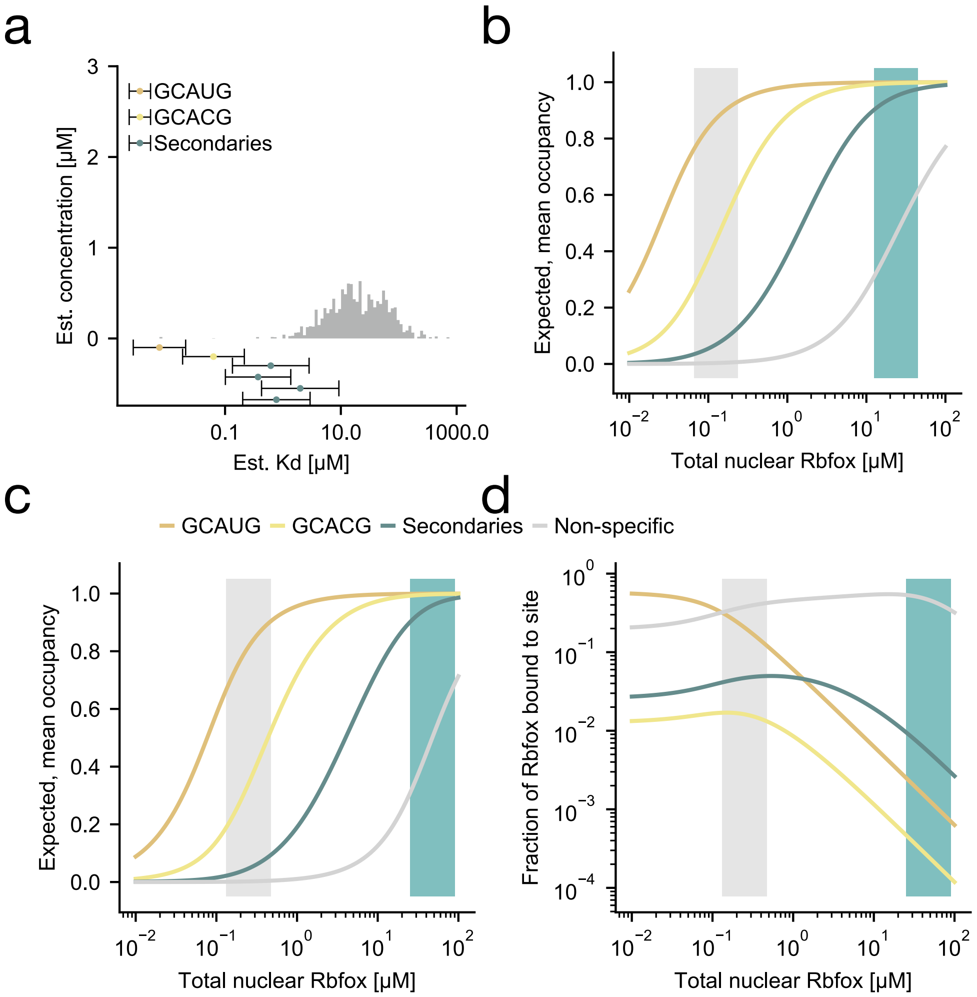

a
b
c
d
GCAUG
GCACG
Secondaries
Non-specific
Est. concentration [μM]
Est. Kd [μM]
0.1
10.0
1000.0
Expected, mean occupancy
Total nuclear Rbfox [μM]
Fraction of Rbfox bound to site

**Figure S10. A model for Rbfox secondary motifs.** The high nuclear mRNA scenario assumes 1,000,000 mRNAs/cell with average half-life time of 3 hours. A corresponding, expression weighted histogram of potential intronic Rbfox binding sites is shown in (a). Motif 5-mers are highlighted in gold (GCAUG), yellow (GCACG), and teal (secondary motifs GCUUG, GAAUG, GUUUG, GUGUG, GUAUG, GCCUG). The low nuclear mRNA model assumes 10x lower mRNA copies/cell and a half-life time of 4 hours, reducing the estimated amount of RNA and the scale of the y-axis to 7.5% of what is shown in (a). Predicted, average Rbfox occupancies on 5-mer motifs as a function of the nuclear Rbfox concentration are shown in (b,c). While 50% occupancy thresholds differ between low (b) and high (c) mRNA scenarios, they robustly predict distinct, partially overlapping Rbfox concentration regimes in which each motif class (colors as in (a)) is responsive to changes in Rbfox expression. Secondary motifs are consistently predicted to be bound at higher Rbfox concentrations than primary motifs. The low mRNA scenario predicts that the fraction of Rbfox bound to secondary motifs surpasses primary motifs at Rbfox levels > 1 µM (d). This is lower than estimates from the high mRNA scenario in main Figure 6 (~14 µM). Non-specific binding is monitored as the predicted, average occupancy of all 5-mers with $K_d$ > 2 x max(secondaries) in (b) and (c), and as the sum of predicted Rbfox bound to these 5-mers in (d).
